# CaMKII-RNMT axis directs activity-dependent control of RNA dynamics in neurons

**DOI:** 10.1101/2025.10.30.685591

**Authors:** Rajaei Almohammed, Shang Liang, Marcus Bage, Lydia Hepburn, Lindsay Davidson, Fiona Haward, Alan Prescott, Haru Yoshikawa, Angus Lamond, Andrei Pisliakov, Victoria H. Cowling

## Abstract

In neurons, mRNAs are transported into axons and dendrites for local translation at synapses. Diverse proteins support the transport of specific mRNAs, however, mechanisms co-ordinating mRNA translocation are unclear. We report that that CaMKII (Ca2+/calmodulin-dependent protein kinase II) is a key regulator of RNA dynamics through activity-dependent phosphorylation of the RNA cap guanine-7 methyltransferase, RNMT. Following RNA cap methylation in the nucleus, RNMT is retained on specific mRNAs. On stimulation, RNMT translocates into the cytoplasm, increasing levels of locally translated mRNAs. In the cytoplasm, CaMKII phosphorylates RNMT Thr317 on the active site, which inhibits methyltransferase activity and targets RNMT for degradation, thus limiting cytoplasmic RNMT function. RNMT mutants protected from CaMKII-dependent degradation increase locally translated, cytoplasmic mRNAs and accelerate neuronal morphogenesis. RNMT Thr317 phospho-mimic co-ordinately decreases locally translated mRNAs in the cytoplasm and reduces differentiation. The CaMKII-RNMT relationship links neuronal activity to the spatial regulation of gene expression in neurons.

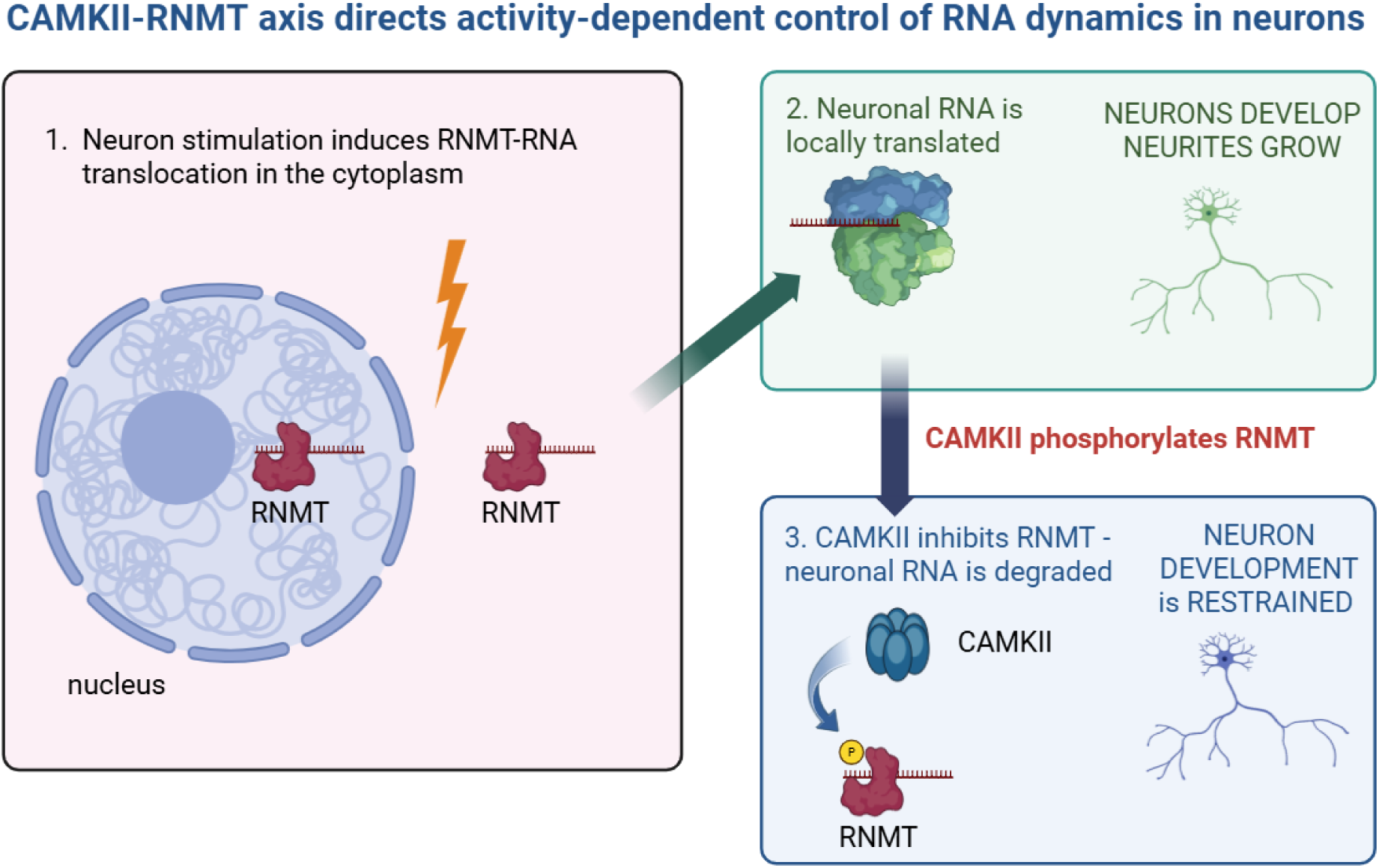

## Introduction

Neurons are specialised cells which transmit information via chemical and electrical signals^1–4^. During differentiation and in response to stimulation, specific mRNAs are translocated long distances along neurites and dendritic spines to be locally translated in synapses^5^. The selective transport and localisation of mRNA involves cis-acting elements interacting with numerous RNA-binding proteins, however, the mechanisms co-ordinating RNA translocation following neuronal activity are undetermined^6,7^. RNA translocation is important in neurons because it contributes to the localised protein remodelling required for neuronal plasticity, including during long-term potentiation and long-term depression^8^. A master mediator of synaptic plasticity is CAMKII (Ca^2+^/calmodulin-dependent protein kinase II) which co-ordinates the function of mechanistic proteins in response to stimulation^9^. The regulation and biological impact of CaMKII in neurons is well described, however its impact on RNA translocation and translation is unclear. Here we report that the RNA cap methyltransferase, RNMT, is redeployed in neurons to have an overarching role in supporting RNA translocation. This redeployment of RNMT in RNA dynamics is regulated by CaMKII.

During transcription, the guanosine cap is added to mRNA and other RNA pol II transcripts^10,11^. RNMT methylates the guanosine cap N-7 position, completing the ^m7^G cap which protects mRNA and recruits proteins involved in RNA processing, export and translation initiation^12,13^. Certain functionally related mRNAs exhibit enhanced responses to RNMT, in a cell-specific manner, with potent impacts in embryonic stem cell differentiation, cell proliferation and T cell activation^14–20^. RNMT is the most abundant and active RNA cap methyltransferase^13,15^. In addition to methylating the RNA cap guanosine on the N-7 position, RNMT binds to RNA directly and via interactions with the activator subunit, RAM^21,22^. RNMT interacts with RNA across the transcriptome, along the full length of transcripts^23^. RNMT-RNA interactions enhance RNA cap methylation but also have a non-catalytic role in transcription^23,24^. Transcription-associated proteins, including the PAF complex, bind to RNMT which increases their recruitment to DNA^23,25^.

Here we report that when neurons are stimulated, RNMT translocates into the cytoplasm where it co-ordinately increases levels of locally translated neuronal mRNAs. This redeployment of RNMT is associated with neurite growth. RNMT function in the cytoplasm is limited by CaMKII-dependent phosphorylation of RNMT T317 in the active site. RNMT T317 phosphorylation reduces methyltransferase activity, co-factor binding and targets RNMT for proteasomal degradation. Expression of RNMT T317 phospho-deficient mutants, which are resistant to CaMKII-dependent degradation, increases locally translated RNA and neurite growth.

## Results

### Depolarisation of neurons induces RNMT translocation from the nucleus to cytoplasm

RNMT is ubiquitously expressed throughout the adult murine brain, with elevated levels in the hippocampus (Figure 1A; confirmed with an independent RNMT antibody, Figure S1A). RNMT is predominantly nuclear in cell lineages investigated to date, including human induced pluripotent stem cells (hiPSC, Figure S1B), consistent with co-transcriptional RNA cap methylation^16,21,26^. In the hippocampal CA1 region, RNMT is present in the nucleus and cytoplasm, in somata and neurites (Figure 1A, S1A). RNMT is cytoplasmic in approximately 30% of primary hippocampal and cortical neurons in culture (16-18 DIV, days *in vitro*, Figure 1B and S2A). When hippocampal and cortical neurons are depolarised (50 mM KCl, 5 mins), RNMT rapidly redistributes from the nucleus to the cytoplasm (30 mins post depolarisation, Figure 1B, 1C and S2A). KCl-dependent depolarisation induces rapid regulation of neuronal gene expression, RNA localisation and RNA-binding proteins shuttling^4,27–31^. In neurons treated with the membrane-permeable Ca^2+^ chelator EGTA (EGTA-AM), RNMT is restricted to the nucleus, indicating that depolarisation-induced translocation is Ca^2+^ influx-dependent (Figure 1B, 1C). In neurons treated with NaCl, an osmolarity control, RNMT is retained in the nucleus (Figure S2B).

**Figure 1.**
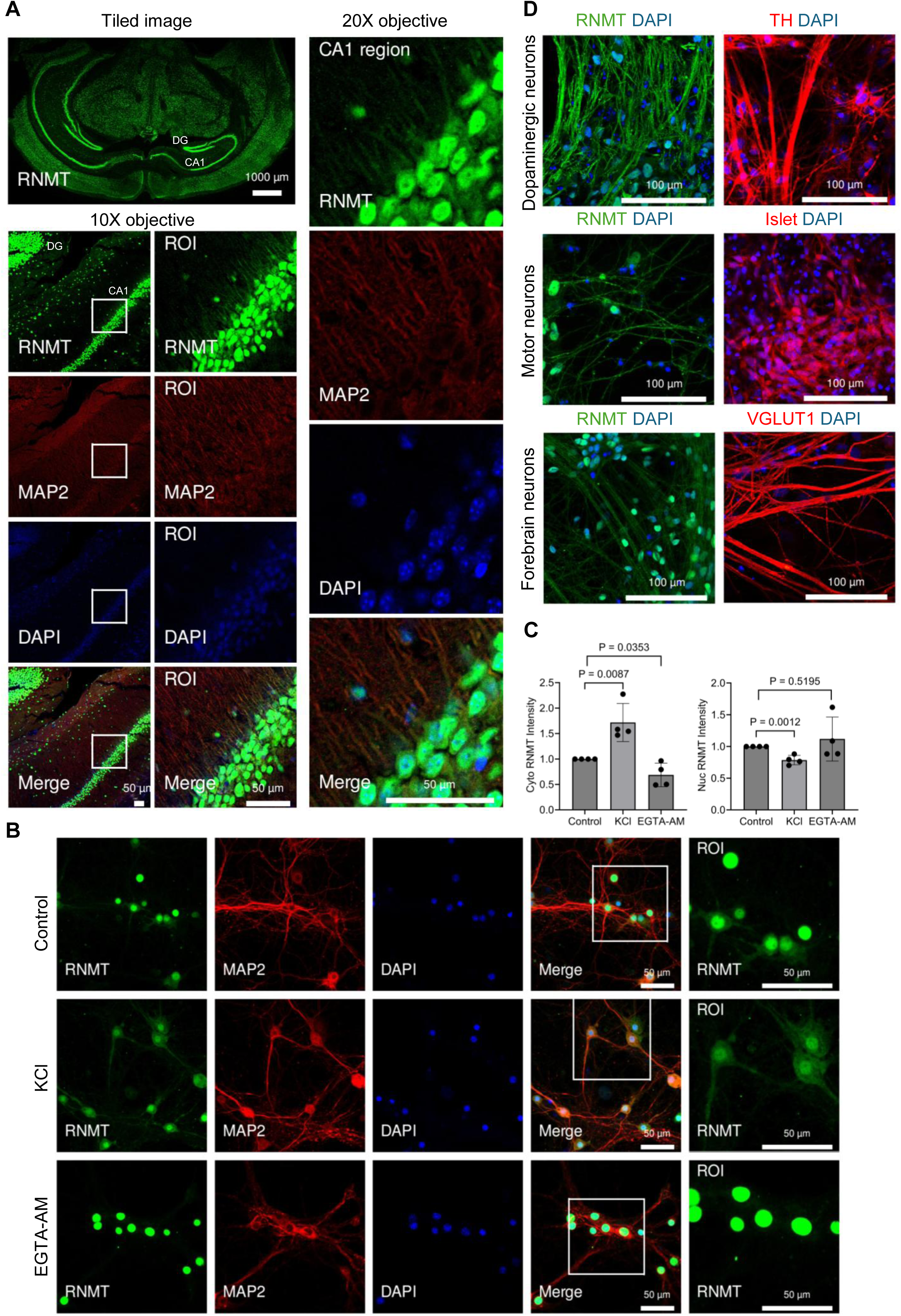
Depolarisation induces RNMT translocation to the cytoplasm. A) Immunofluorescence (IF) analysis of adult mouse brain. Dentate gyrus (DG) and cornu ammonis (CA1) regions are labelled. Sections were stained for RNMT (sheep, green), DAPI (4′,6-diamidino-2-phenylindole) stain (blue), and dendritic marker MAP2 (red). Cytoplasmic RNMT is observed at the end of the hippocampus CA1 region. RNMT staining performed using an alternative rabbit antibody in Figure S1A. B) RNMT staining in primary hippocampal neurons (DIV16). RNMT cytoplasmic localisation increases following neuron depolarisation with 50 mM KCl, compared to DMSO treated controls. RNMT cytoplasmic localisation decreases on ion chelation with 50 µM EGTA-AM. RNMT (green), MAP2 (red), DAPI (blue). See Figure S2A for RNMT IF in primary cortical neurons. C) Quantification of relative cytoplasmic and nuclear RNMT integrated intensity in primary neurons compared to control sample. Error bars represent the mean ± standard deviation (SD) of four independent samples (dots). Results were analysed using unpaired student t-test. P values relative to control indicated. D) RNMT staining in hiPSC-derived neurons: Dopaminergic neurons (DA), Motor neurons and Forebrain neurons. Cells were stained for RNMT (green) and DAPI (blue). Neuron lineage markers are red: DA neurons marker, Tyrosine hydroxylase (TH); Motor neurons, Islet; and Forebrain neurons, VGLUT1.

RNMT is present in the nucleus and cytoplasm in hiPSC-derived neurons; in dopaminergic neurons (DA, TH-positive), motor neurons (Islet-positive), and forebrain neurons (VGLUT1-positive, Figure 1D). Of note, in hiPSC-derived neurons RNMT is cytoplasmic without depolarisation (Figure 1D). Neurotrophins, including BDNF (Brain-derived neurotrophic factor), supplemented to promote differentiation in culture modulate synaptic efficacy and neuronal activity, which may promote RNMT translocation^32,33^. When expressed in DA neurons and primary murine neurons, GFP-RNMT is also in the nucleus and cytoplasm (Figure S3A and B), whereas in parental hiPSC, GFP-RNMT (and endogenous RNMT) is primarily in the nucleus (Figure S1B). The pre and post-synaptic compartments of neurons contain unique proteins and transcripts^34,35^. In DA neurons, equivalent levels of RNMT are present in MAP2 and Synapsin I-positive neurites, consistent with localisation in dendrites and axons (Figure S3C). Taken together, these data indicate that RNMT may have a function in the cytoplasm of neurons.

### RNMT interacts with CaMKII and mRNAs in neurons

To determine its function in neurons, RNMT-interacting mRNAs and proteins were identified. Single-end enhanced UV cross-linking and immunoprecipitation (seCLIP) analysis was performed in primary murine neurons to identify mRNAs which interact with RNMT^36^ (Figure 2A). RNMT interacts with mRNA at the transcription start site, adjacent to the guanosine cap, but also across the transcript body (Figure 2A, Table S1). Specific mRNAs are enriched on RNMT in neurons (see below). RNMT is complexed with its co-activator RAM in cell lines investigated to date, including embryonic stem cells (ESC)^26,37^. Gel filtration analysis revealed that RNMT is present in larger complexes in the murine brain compared to ESC, indicating interactions with additional or alternative proteins to RAM (Figure 2B). RNMT complexes were purified from mouse brain extracts and interacting proteins were identified by mass spectrometry (MS, Table S2). Established RNMT interacting proteins were detected, including RAM^21^. Cytoplasmic proteins, including Ca^2+^/calmodulin-dependent protein kinase II (CaMKII) subunits, were identified in RNMT complexes (Figure 2C, Table S2, S3).

**Figure 2.**
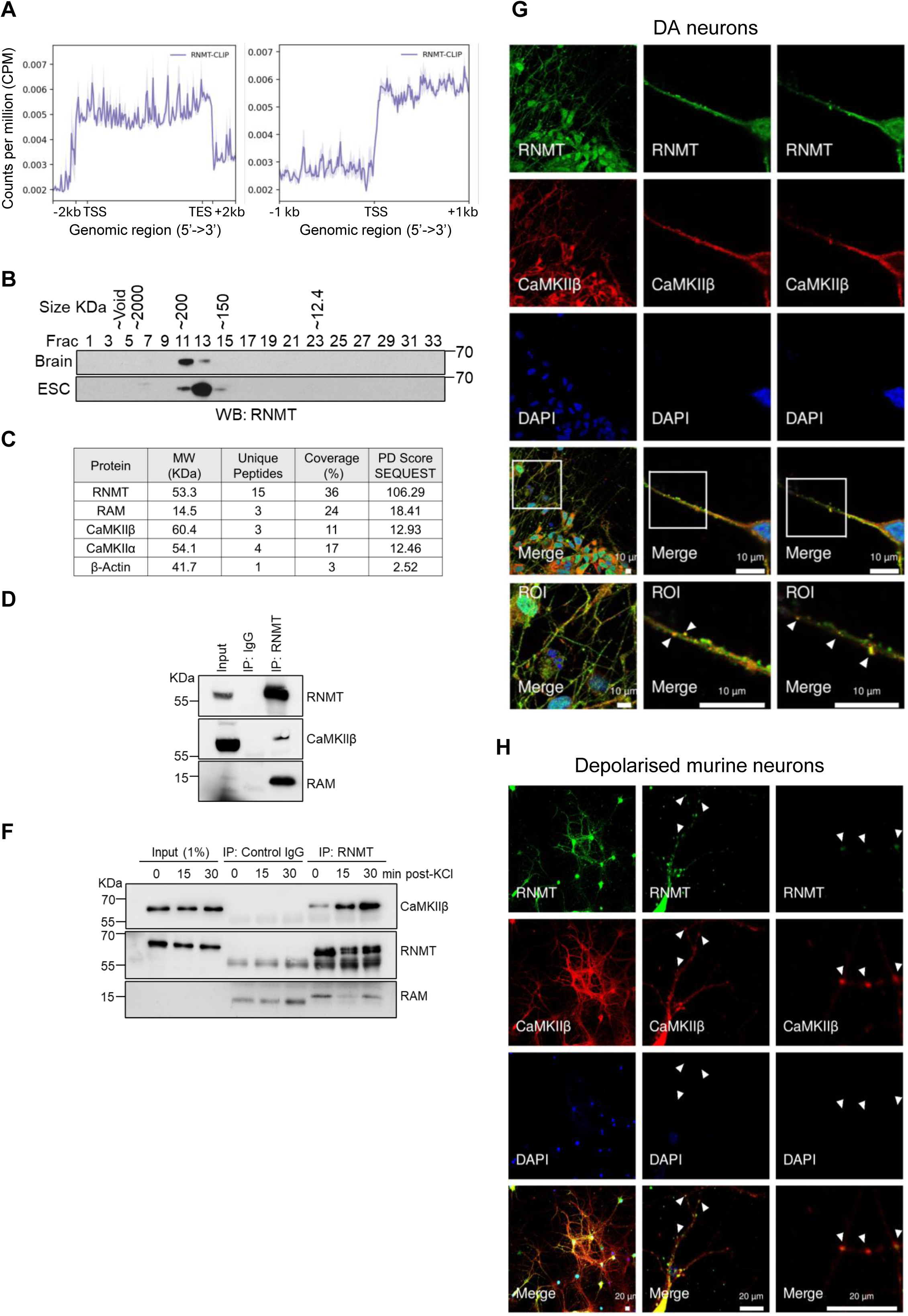
RNMT interacts with CaMKII in neurons. A) seCLIP analysis of RNMT-RNA interaction in murine neurons. RNA was UV-cross linked to protein in neurons. RNMT was immunoprecipitated, RNA extracted and analysed by RNA sequencing. Reads mapped to the genome are presented. Transcription start site (TSS) and end site (TSE) indicated. B) Mouse brain and ESC extracts were resolved by gel filtration on Superdex S200 column. Fractions were analysed for RNMT expression by western blotting. The migration of standards and the void volume are indicated. C) RNMT was immunoprecipitated from adult mouse brain extracts and complexes analysed by mass spectrometry. For RNMT, RAM, CaMKIIα, CaMKIIβ and β-Actin unique peptides, coverage, and Proteome Discoverer (PD) score are given. D) In adult mouse brain extract, RNMT was immunoprecipitated using sheep anti-RNMT antibody. RNMT, RAM, and CaMKIIβ were detected in the immunoprecipitates. F) Primary neurons were incubated with 50mM KCl for 5 mins then left to recover for 15 or 30 minutes. RNMT was immunoprecipitated. Increasing amount of CaMKIIβ was detected in complex with RNMT following KCl activation. RNMT activating subunit RAM was used as a control. G, H) RNMT and CaMKIIβ detected by immunofluorescence in G) hiPSC-derived DA neurons and H) depolarised primary hippocampal neurons. RNMT (green) and CaMKIIβ (red) co-localisation in neurites is marked by white arrows. DAPI nuclear staining (blue).

CaMKII is an abundant, 12-subunit kinase found predominantly in the cytoplasm of neuronal cells. In brain tissue, CaMKIIα homomers and CaMKIIα–CaMKIIβ heterodimers are the most abundant CaMKII complexes^39,40^. Following neuron stimulation, CaMKII is activated by Ca^2+^ influx. Through phosphorylation of its substrates, CaMKII co-ordinates responses to depolarisation and is crucial for hippocampal long-term potentiation^41^. CaMKII phosphorylates key substrates to mediate the response to stimulation, including AMPA-type glutamate receptors^42^. However, little is known about the impact of CaMKII on gene expression, with the exception of regulation of CPEB impacting on local translation^43,44^.

RNMT interaction with CaMKIIβ was verified by western blot analysis of RNMT immunoprecipitates from brain extracts (Figure 2D). Following depolarisation, increasing amount of CaMKII is purified with RNMT from murine neurons, consistent with RNMT translocation into the cytoplasm (Figure 1B, 1C, 2F, confirmed by MS analysis, Table S3, Figure S4A, B).

To map RNMT and CaMKII interaction, GFP-CaMKIIα was expressed with HA-RNMT WT and mutants in HEK293 cells. GFP-CaMKIIα immunoprecipitated via the GFP tag interacts with full length HA-RNMT and HA-RNMT catalytic domain (HA-RNMTcat, 121-476), but not with the N terminal regulatory region (HA-RNMT-N, 1-120). Conversely, HA-RNMT and HA-RNMTcat immunoprecipitated via the HA tag bind to GFP-CaMKIIα (Figure S4C). CaMKIIβ and RNMT co-localise in the neurites of hiPSC-derived DA neurons and depolarised primary murine neurons (Figure 2G, 2H). In neurites RNMT is predominantly diffusely distributed, however, RNMT colocalises with CaMKII in puncta, consistent with postsynaptic CaMKII localisation^45^.

### CaMKII phosphorylates RNMT on Threonine 317

CaMKII predominantly impacts on cellular functions by phosphorylating substrates, thus altering their activity, interactions or stability^9^. CaMKII was observed to phosphorylate RNMT *in vitro* (Figure 3A, 3B and S5A). Recombinant GST-CaMKIIα (activated by incubation with calmodulin and ATP), was incubated with recombinant His-RNMT and γ^32^P-ATP. A ^32^P-labelled band consistent with RNMT phosphorylation was detected after 5 minutes (Figure 3A). CaMKIIα also autophosphorylates (Figure 3A and S5A)^46,47^. The catalytic domain of RNMT (165-476, RNMTcat), which interacts with CaMKII, is phosphorylated by CaMKIIα (Figure S4C, 3B). CREB, an established CaMKIIα substrate, was used as a control^48^. To map phosphorylation sites, recombinant RNMT was incubated with CaMKIIα and analysed by MS. Multiple phosphorylation sites were identified (Figure 3C, S5C, Table S4). To focus on physiologically relevant phosphorylation sites, RNMT purified from murine brain and ESC was analysed. Phosphorylated T317, T318 and S321 were detected in RNMT purified from brain extracts but not in ESC extracts (Figure 3C and Table S4). RNMT T317 and S321 are also phosphorylated by CaMKIIα *in vitro* (Figure 3C, Table S4). Phosphorylation of RNMT T317, T318 and S321 is also observed in the promyelocytic leukaemia cell line, HL-60^49^.

**Figure 3.**
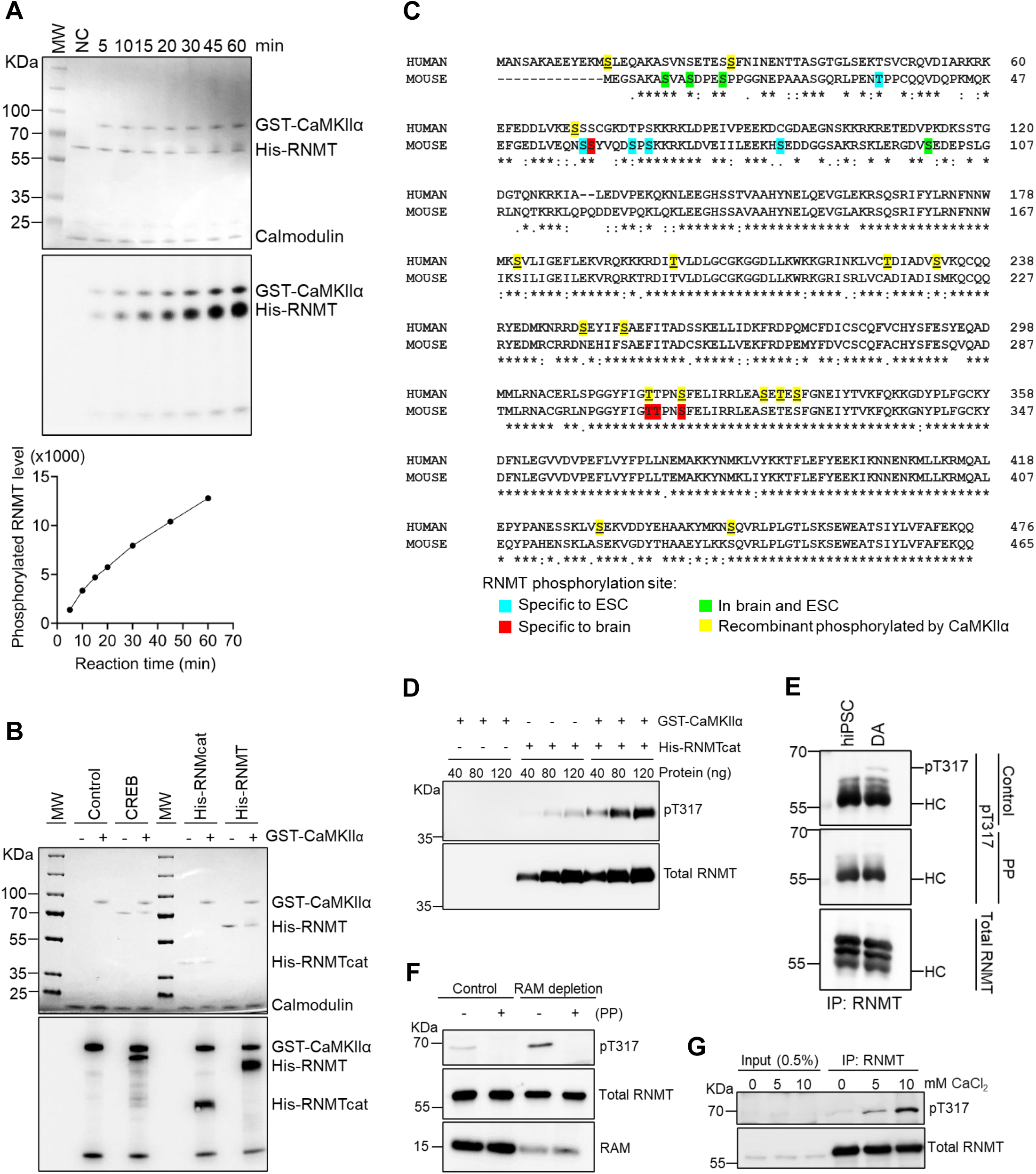
RNMT is phosphorylated on T317 by CaMKII. A) Recombinant His-RNMT was incubated with activated GST-CaMKIIα and ^32^P-γATP for 60 minutes. Proteins were visualised on a Coomassie stained gel (upper panel). Phosphorylated RNMT bands were detected by phospho-imaging (bottom panel). Molecular weight (MW) marker and no kinase negative control (NC) indicated. Quantitation of phosphorylation presented. B) Recombinant RNMT WT and catalytic domain (His-RNMTcat, 165-476aa) was incubated with GST-CaMKIIα and ^32^P-γATP. CREB was used as a positive control. Proteins were visualised in a Coomassie stained SDS-PAGE gel (Upper panel). Phosphorylated proteins detected by phospho-imaging (bottom panel). C) Mass spectrometry analysis. RNMT phosphorylation sites identified in mouse embryonic stem cell (ESC) lysate, mouse brain lysate and recombinant RNMT *in vitro* phosphorylated by CaMKII. D) An antibody was raised in sheep against an RNMT pT317 peptide, 312 GYFIG**T**TPNSF 322. The RNMT pT317 antibody was used in a western blot to detect recombinant RNMT phosphorylated by CaMKII. (E-G) RNMT was immunoprecipitated and endogenous pT317 and total RNMT were detected by western blot: E) hiPS cells and hiPSC-derived DA neurons. Lambda protein phosphatase (PP) or control buffer was incubated with pT317 membranes. F) mouse brain lysate, RAM immuno-depleted prior to RNMT IP. Lambda protein phosphatase (PP) incubated with RNMT IPs, as indicated. G) Brain lysate incubated with CaCl_2_.

The function of RNMT phosphorylation was investigated, focusing on Threonine 317 (T317) which is in the active site^24^. Polyclonal antibodies were raised against human RNMT 312 GYFIG**T**TPNSF 322, phosphorylated on T317. By western blot, the RNMT pT317 antibody detects recombinant RNMT phosphorylated by CaMKIIα (Figure 3D). GST-CaMKIIα phosphorylates RNMT monomer but not the RNMT-RAM complex (Figure S5B). Although RNMT T317 is distal to the residues involved in RAM interaction, RAM binding alters RNMT dynamics which may make T317 inaccessible to CaMKIIα^24,50^. The RNMT pT317 antibody also detects a band consistent with phosphorylated GST-CaMKIIα, likely due to the pT317 antibody recognising similar CaMKII substrate motifs on CaMKII and RNMT (Figure S5B). In western blot analysis of RNMT IPs from brain lysates, a band is detected by the RNMT pT317 antibody which migrates more slowly than total RNMT and decreases with phosphatase treatment, consistent with detection of RNMT phospho-T317 (Figure 3F). On depletion of RNMT-RAM complexes from cell extracts with an anti-RAM antibody, detection of RNMT pT317 increases, consistent with RNMT and not the RNMT-RAM complex being a CaMKII substrate (Figure 3F, S5B).

The RNMT pT317 antibody was used to detect RNMT phosphorylation in cells. RNMT pT317 is detected in hiPSC-derived DA neurons but not the parental hiPS cells, consistent with neuron-specific phosphorylation (Figure 3E). CaMKII is activated by calcium signalling. Increased RNMT pT317 is detected in brain homogenates incubated with CaCl_2_, consistent with CaMKII-dependent phosphorylation of RNMT (Figure 3G).

### CaMKII-dependent phosphorylation supresses RNMT catalytic activity

Neuron stimulation results in RNMT translocation from the nucleus to the cytoplasm (Figure 1), where it interacts with CaMKII (Figure 2), and is phosphorylated by CaMKII (Figure 3). We investigated the molecular impact of RNMT T317 phosphorylation, focusing on substrate binding and guanosine cap methylation. In the RNMT active site, the methyl donor s-adenosyl methionine (SAM) and the guanosine cap substrate are positioned to facilitate methyl transfer^24^. Recombinant RNMT was phosphorylated by incubation with CaMKIIα and ATP, and the impact on methyltransferase activity was measured. A ^32^P-capped transcript was incubated with RNMT and SAM. RNA cap methylation is reduced ∼3-fold when RNMT is phosphorylated by CaMKII (Figure 4A).

**Figure 4.**
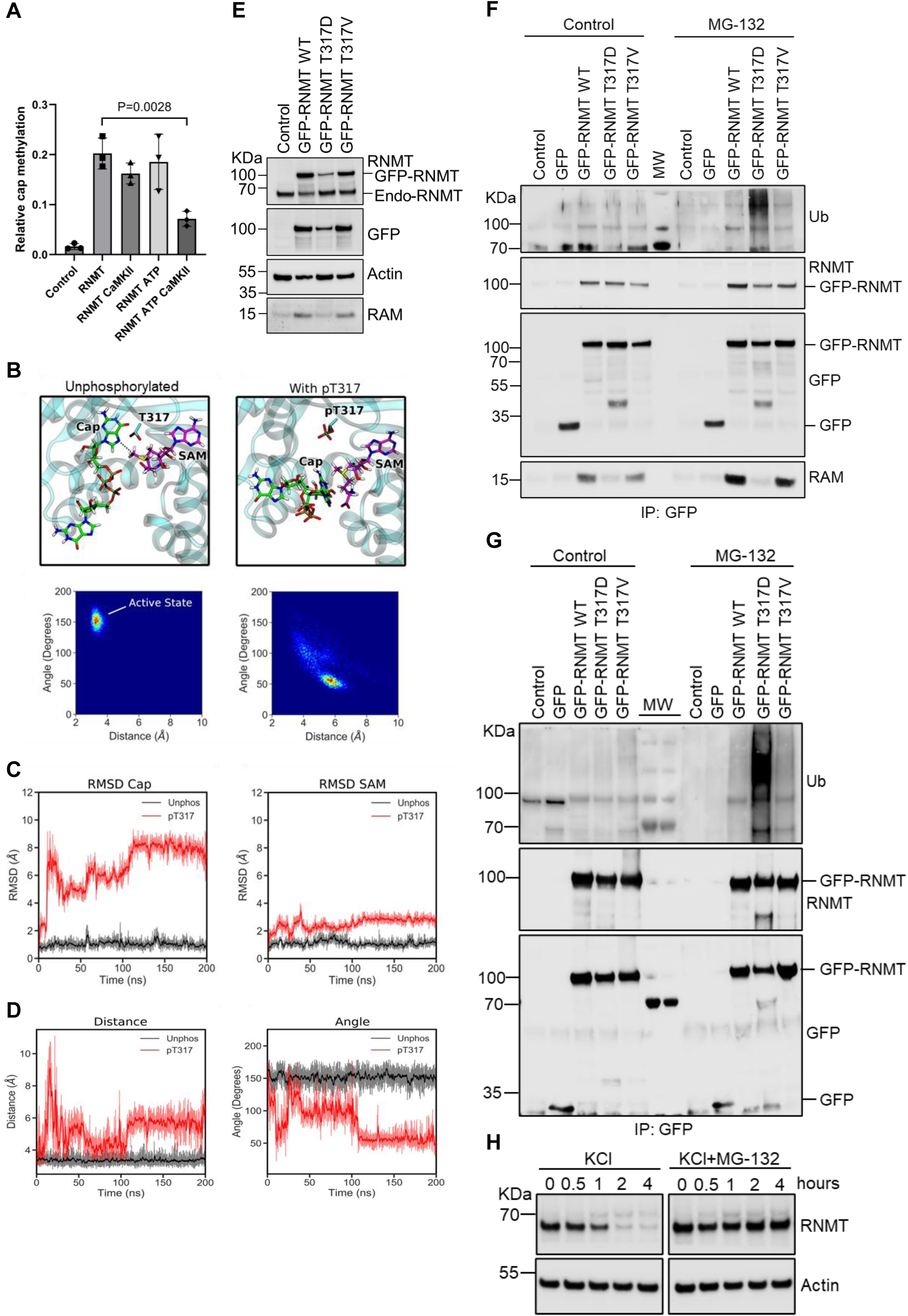
RNMT T317 phosphorylation inhibits its function. A) Recombinant RNMT was incubated +/-CaMKII and ATP for 90 minutes. The RNA cap methyltransferase assay was performed. Data from 3 independent experiments are presented. Error bars represent the mean ± SD. Results were analysed using unpaired student t-test. B) Molecular Dynamics simulations of RNMT, s-adenosyl methionine (SAM) and unmethylated cap (GpppG). Substrate coordination in the unphosphorylated and T317 phosphorylated states over 200ns. Top panel, representative snapshots showing the most occupied conformation of the RNMT substrates in the unphosphorylated and pT317 states during the MD simulations (Figure S7C, Systems 1 and 2). Bottom panel, heatmaps displaying the distance (cap G0 N7 – SAM methyl group) and angle (cap G0 N9 – G0 N7 – SAM methyl group) of the RNMT reactive groups during MD simulations of RNMT in the unphosphorylated and pT317 states (Systems 1 and 2). The active state (pre-catalysis) coordination of the ligands corresponds to a distance of ∼3.5 Å and ∼150 degrees. C) Stability of the RNMT substrates in the unphosphorylated and T317 phosphorylated states. Time-evolution RMSDs of the cap (G0) and SAM over the duration of MD simulations of RNMT in the unphosphorylated (black) and pT317 (red) states (Systems 1 and 2). D) Quantification of substrate coordination in RNMT in the unphosphorylated and T317 phosphorylated states. Time-evolution of the distance (cap G0 N7 – SAM methyl group) and angle (cap G0 N9 – G0 N7 – SAM methyl group) of the substrate reactive groups during MD simulations of RNMT in the unphosphorylated (black) and pT317 (red) states (systems 1 and 2). E) Expression levels of GFP-RNMT WT, T317V and T317D in HEK293 cells analysed by western blots. F) HEK293 cells expressing RNMT T317 mutants were treated with 20 μM MG-132 for 3 hours. GFP-RNMT T317 mutants and GFP control were immunoprecipitated via the GFP tag (GFP-RNMT immunoprecipitated from each cell line was balanced by altering the quantity of lysate added to each GFP-IP, see Figure S8). Western blot performed to detect GFP-RNMT, GFP, RAM and ubiquitinated protein. See Figure S8A and B for establishing RNMT T317 mutants in HEK293 cells. G) SH-SY5Y cells expressing RNMT T317 mutants were treated with 20 μM MG-132 for 3 hours. GFP immunoprecipitation and western blot analysis were performed to detect RNMT, GFP and ubiquitinated protein. H) Primary murine hippocampal neurons were treated with 50 mM KCl with and without 20 μM MG-132. RNMT expression was detected by western blots. Actin was used as a loading control.

To understand the mechanism by which RNMT T317 phosphorylation reduces methyltransferase activity, molecular dynamics (MD) simulations were performed to model SAM and guanosine cap binding in the active site. An electrostatic surface representation shows that T317 phosphorylation significantly alters the electrostatic environment of the active site (Figure S6A). The cap binding site is substantially more negatively charged when T317 is phosphorylated, with smaller changes in electrostatics in the SAM binding site. In simulations of substrate-bound, catalytically primed RNMT, with T317 unphosphorylated, substrates remain in the active conformation. This can be quantified by measuring the distance (cap G0 N7 – SAM methyl group) and angle (cap G0 N9 – G0 N7 – SAM methyl group) between the reactive groups where the catalytic distance remains at ∼3.5 Å and the catalytic angle remains at ∼150 degrees (Figure 4B). In contrast, phosphorylation of T317 resulted in significant destabilisation of cap and SAM in the active site. After 5 ns, the cap was displaced from its binding pocket and did not rebind, and the catalytic distance and angle became more variable (Figure 4C, 4D). In the most occupied state, the catalytic distance increases to ∼6 Å and the catalytic angle decreases to ∼50 degrees.

RNMT T317 lies adjacent to F285, which pi-stacks the G0 (unmethylated) cap^50^. In MD simulations, phosphorylation of T317 causes conformational changes in residues that coordinate the cap, with SAM coordination less affected (Figure 4B and S6B). As the F285 sidechain is hydrophobic, it is repulsed when there is a negatively charged phosphate on T317, causing it to rotate away into a conformation unsuitable for pi-stacking with the cap. This conformational change destabilises the coordination of the cap, resulting in dissociation from the cap-binding pocket. MD simulations of apo-RNMT are similar to the substrate-bound systems, with significant conformational change in the cap-binding site on T317 phosphorylation (Figure S6C). Thus, T317 phosphorylation alters the RNMT active site, reducing affinity for the cap, correlating with reduced methyltransferase activity.

### The phospho-mimic T317D supresses RNMT activity and reduces stability

To determine the impact of RNMT phosphorylation in neurons, MD simulations were used to identify amino acid substitutions which effectively mimic T317 phosphorylation status. T317D or T317E were assessed to be suitable mimics of RNMT T317 phosphorylation (Figure S7A). T317V (not T317A), replicates the behaviour of unphosphorylated RNMT (Figure S7B). A summary of simulations performed is presented in Figure S7C. GFP-RNMT WT, phospho-defective T317V and phospho-mimic T317D mutants were expressed in HEK293 cells. GFP-RNMT IPs were analysed by western blot (Figure S7D) and used in the RNA cap methyltransferase assay (Figure S7E). The WT and T317V mutant methylate the guanosine cap (produce m7GpppG) equivalently. Consistent with being a phospho-mimic, RNMT T317D is inactive. Catalytic-dead mutant D203A was used as a control ^51^.

To investigate the cellular function of RNMT T317 phosphorylation, HEK293 cell lines were generated to express GFP-RNMT WT, T317D and T317V. GFP-RNMT T317D is expressed at a reduced level compared to WT and T317V mutant (Figure 4E). When incubated with the proteosome inhibitor, MG-132, high molecular weight ubiquitinated protein is detected in the GFP-RNMT IPs, and this is enhanced in the T317D but not T317V mutants (Figure 4F). (In this analysis 5-fold more extract was analysed from T317D-expressing cells compared to other cell lines, to equalise GFP-RNMT levels across the panel.) This data is consistent with the phospho-mimic, RNMT T317D, having enhanced ubiquitin-dependent proteasomal degradation. Similar observations were made on expression of GFP-RNMT WT, T317D and T317V in SH-SY5Y cells, a neuroblastoma cell line (Figure 4G). RNMT stability is dependent on binding of the co-factor RAM^21,52^. T317D is defective for RAM binding, which is likely to contribute to its instability (Figure 4E, 4F, S7D).

To investigate if the stability of endogenous RNMT is also repressed in correlation with CaMKII-dependent phosphorylation, primary murine neurons were depolarised with KCl (Figure 4H). Within 2 hours of depolarisation, RNMT levels are significantly repressed. Consistent with proteasome-mediated degradation, incubation with MG-132 prevents RNMT repression in depolarised neurons.

RNMT binds to the guanosine cap but also to RNA directly^21,23^. The impact of RNMT phosphorylation on RNA binding was modelled by expressing GFP-RNMT T317 mutants in HEK293 cells and performing GFP seCLIP^36^. HEK293 cells were utilised because RNMT-RNA interactions are difficult to capture in iPSC-derived neurons due to technical limitations. Since the T317D mutant is expressed at a reduced level compared to the WT protein or T317V, the amount of GFP-RNMT immunoprecipitated from each cell line was balanced by altering the quantity of lysate added to each GFP-IP (Figure S8). RNA binds to RNMT WT, T317V and T317D, detected as a smear of RNMT-RNA complexes in gel electrophoresis (Figure 5A).

**Figure 5.**
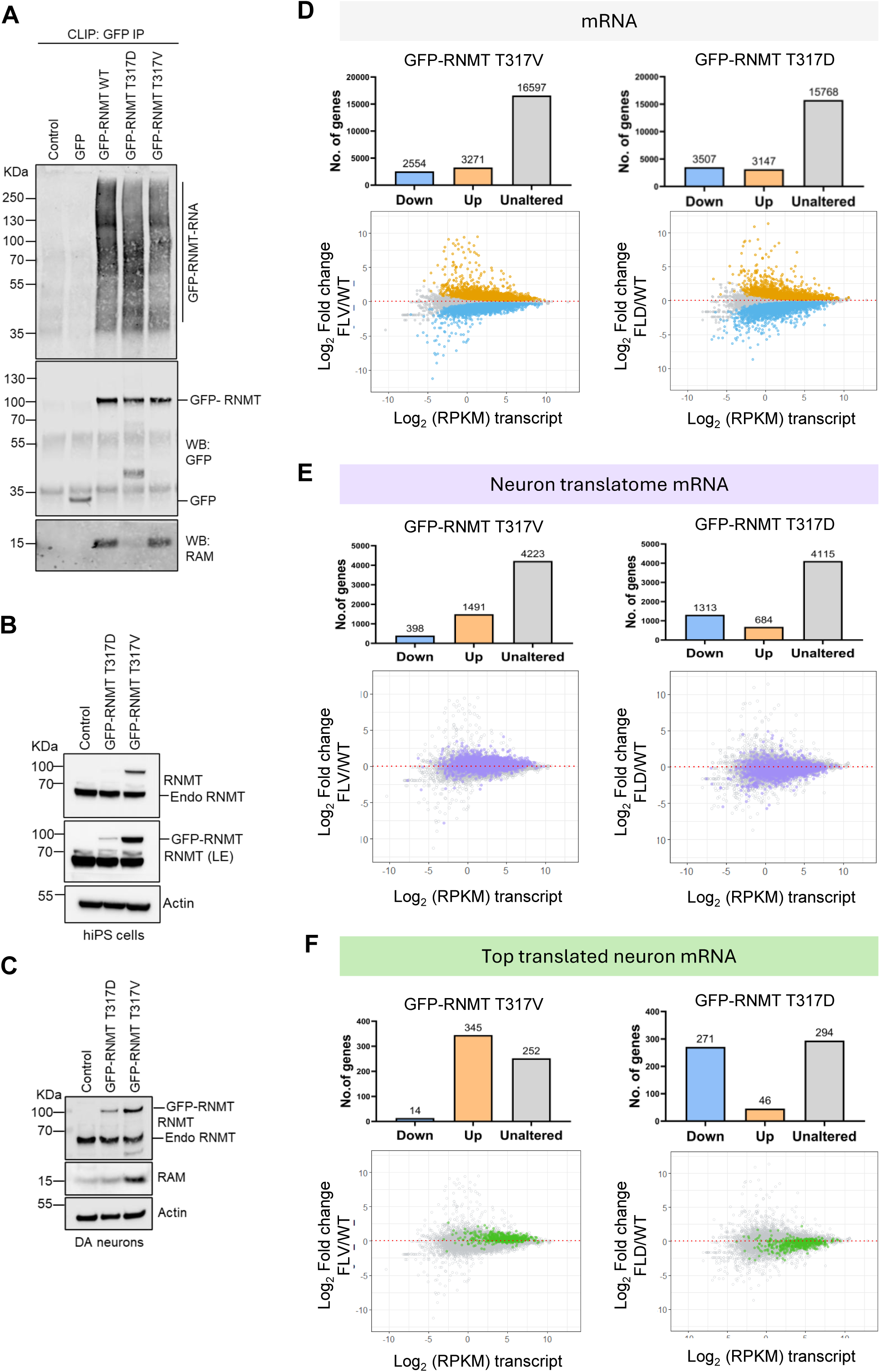
RNMT regulates a subset of locally translated mRNA. A) seCLIP analysis of RNMT-RNA interactions in HEK293 cell lines. RNA was UV-cross linked to protein in cells. GFP, GFP-RNMT WT, T317D and T317V were immunoprecipitated via the GFP tag. GFP-RNMT immunoprecipitated from each cell line was balanced by altering the quantity of lysate added to each GFP-IP, see Figure S8. RNA was fragmented, ligated to fluorescent adaptors, resolved by gel electrophoresis and visualised by LI-COR imaging system (upper panel). Immunoprecipitated GFP-RNMT mutants, GFP control and RAM were analysed by western blotting (lower panel). B) Western blot analysis of GFP-RNMT T317D and T317V expression in hiPSC cells (ChiPSC4). Longer exposure (LE) of RNMT blot presented. Actin is a loading control. C) Western blot analysis of GFP-RNMT T317D and T317V expression in hiPSC-derived DA neurons. RAM and actin blots presented. D-F) RNA sequencing analysis of DA neurons. For genes, scatterplot of transcript level change (Log_2_ fold change) and transcript abundance (Log_2_ RPKM, reads per kilobase per million mapped reads). Differentially expressed transcripts were identified with false discovery rate (FDR)<0.05. D) Downregulated (blue), unaltered and upregulated (orange) transcripts are indicated. E) translated neuronal transcripts identified in ^55^ are highlighted (Neuron translatome mRNA, purple). F) Top 10% of the translated transcripts indicated (Top translated neuron mRNA, green). Bar charts show number of downregulated, unaltered and upregulated transcripts for each cell line.

### RNMT regulates a subset of locally translated mRNA

Since CaMKII co-ordinates mechanisms during neuron development and stimulation, we investigated the impact of CaMKII phosphorylation of RNMT in neurons using the T317 mutants. hiPSC lines expressing either GFP-RNMT-T317D or GFP-RNMT-T317V were made using a transposon system^53^, Figure 5B). The RNMT T317D mutant was expressed at a lower level than the T317V mutant, as observed in other cell lines (Figure 4E and S8). All generated hiPS cell lines maintained the ability to form embryoid bodies. Similar to endogenous RNMT, GFP-RNMT WT, T317D and T317V are predominantly nuclear in hiPSC (Figure S1B, S9A). The hiPSC lines were differentiated to DA neurons^54^ (Figure 5C, S9B). Neural conversion was induced by exposing embryoid body (EB) to dual SMAD inhibition prior to the derivation of neurons. As observed with endogenous RNMT and GFP-RNMT, the GFP-RNMT mutants T317D and T317V are expressed in the nucleus and cytoplasm of DA neurons (Figure 1D, S3A, S9C). GFP-RNMT WT, T317V and T317D transiently expressed in primary murine neurons are also present in the nucleus and cytoplasm (Figure 1B, S3B and S9D).

RNMT binds to RNA in the nucleus of neurons (Figure 2A). RNMT is translocated into the cytoplasm on depolarisation (Figure 1B, 1C, S2), where it interacts with CaMKII (Figure 2), and is phosphorylated and degraded (Figure 3 and 4). The impact of RNMT on mRNA present in the cytoplasm was investigated. RNMT T317V was used to model the impact of RNMT transport into the cytoplasm, without CaMKII-dependent degradation. The phospho-mimic RNMT T317D mutant was used to model unrestrained phosphorylation-dependent degradation of RNMT. Cytoplasmic RNA was extracted from DA neurons expressing GFP-RNMT T317D, T317V or wild-type cells (control) and RNA sequencing analysis was performed (Figure 5D-F, Table S5). In DA neurons expressing GFP-RNMT T317V, of 22422 transcripts passing detection thresholds, 3271 were upregulated and 2554 were downregulated, compared to control DA neurons. In DA neurons expressing RNMT T317D, 3507 transcripts were downregulated and 3147 were upregulated compared to control cells (Figure 5D).

Functionally related mRNAs were co-regulated in response to RNMT. Synapse-related mRNAs were enriched in the 20% most upregulated mRNAs in DA neurons expressing T371V and enriched in the 20% most repressed mRNAs in DA neurons expressing T317D (Gene Ontology analysis, Figure S9E). Given the enrichment for synaptic function in RNMT-responsive mRNAs, a published study of the transcriptome and translatome in different neuronal compartments was utilised to provide datasets for further analysis^55^. In this study, ribosome sequencing was used to determine the translation level for transcripts in cell bodies (somata) and synaptic regions (neuropil) of excitatory neurons in rodent hippocampus (data set termed “neuron translatome mRNA”, Table S5 column AF: 6112 transcripts)^55^. From the neuron translatome mRNA dataset, over 20% of transcripts were upregulated in the cytoplasm of DA neurons expressing RNMT T317V and over 20% transcripts were downregulated in DA neurons expressing RNMT T317D, compared to controls (Table S5, Figure 5E). A further dataset was created of the top 10% translated neuronal mRNAs from the published neuronal translation study^55^, (termed “top translated neuron mRNA”, Table S5, column AE: 611 mRNAs). 345 mRNAs from the top translated neuron mRNA group were upregulated in neurons expressing T317V compared to control cells, including 44 that have increased translation levels in neuropil. In neurons expressing RNMT T317D, 271 mRNAs of the top translated neuron mRNA were repressed, including 42 that have increased translation levels in neuropil (Figure 5F).

In summary, there is an enrichment of the highly translated, neuronal mRNAs in the transcripts upregulated in response to RNMT T317V, which is protected from CaMKII-dependent degradation. There is an enrichment in highly translated, neuronal mRNAs in transcripts down regulated by the RNMT phospho-mimic T317D, which has enhanced instability, mimicking CaMKII-independent degradation.

### RNMT promotes neurite growth in DA neurons

The increased levels of synapse-related mRNAs in the cytoplasm of DA neurons expressing RNMT T317V could result from several mechanisms. To determine whether the mRNAs increased in the cytoplasm are also elevated in the nucleus, RT-PCR was performed on RNA isolated from sub-cellular fractions. The expression of locally translated mRNAs SYN1, CAMK2B, BDNF and GRM7 was assessed (Figure 6A). In cytoplasmic mRNA extracted from DA neurons expressing T317V, CAMK2B, BDNF and GRM7 mRNAs were increased, whereas in DA neurons expressing RNMT T317D, SYN1, CAMK2B, BDNF and GRM7 mRNAs were repressed. This confirms results from RNA sequencing data (Figure 5E-F, Table S5). In the nucleus, only SYN1 mRNA was upregulated in response to RNMT T317V and repressed in response to RNMT T317D. EIF4A1 and EIF4A2 mRNA, analysed as controls, were at equivalent levels in all cell lines and fractions. These data are consistent with RNMT controlling RNA export or stability in the cytoplasm. In correlation with changes in mRNA level, Syn1, CaMKIIβ, and the neuronal marker Tuj1 were repressed at the protein level in response to expression of RNMT T317D compared to either T317V or control (Figure 6B).

**Figure 6.**
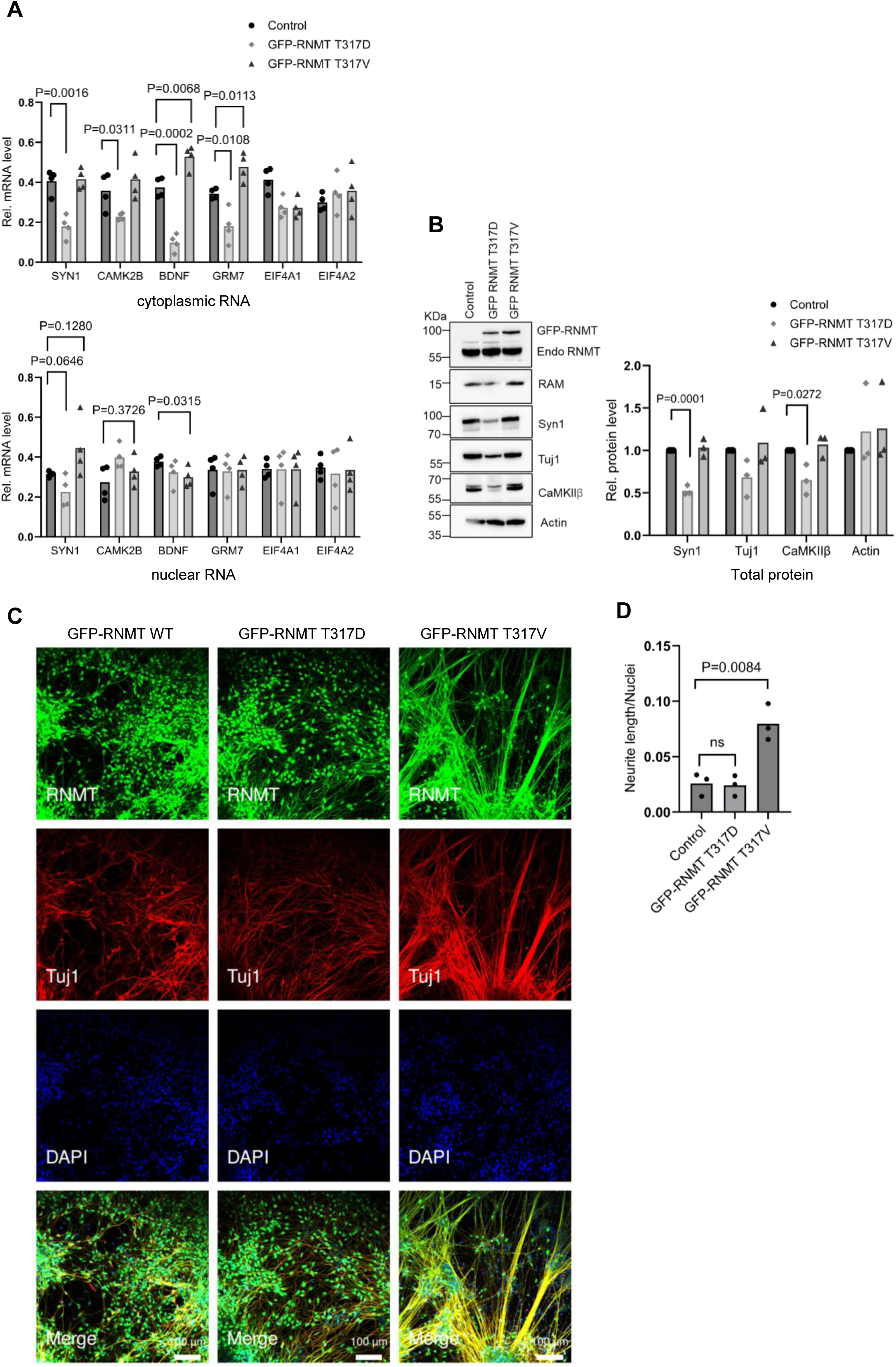
RNMT promotes neurite growth in DA neurons. A) RNA from cytoplasmic (top) and nuclear (bottom) fractions of hiPSC-derived DA neurons, expressing RNMT-GFP-T317V and RNMT-GFP-T317D or control. Transcripts were detected by RT-PCR, normalised to GAPDH. Data represent the mean ± SD of 4 independent samples analysed using unpaired t test. P-values relative to control indicated. B) hiPSC-DA neuron lines were analysed for protein levels by western blotting (a representative panel is shown). Actin was used as a loading control. Protein levels were quantified from three independent experiments, analysed using unpaired t test. P-values indicated. (C) hiPSC-DA neurons expressing GFP-RNMT T317D, GFP-RNMT T317VD or control. RNMT (green), Tuj1 (Red), and DAPI nuclear staining (blue). D) Neurite length was determined by Incucyte Neurotrack and normalised to number of nuclei. Data are representative of three independent experiments, analysed using unpaired t test. P-values indicated.

CaMKII plays a pivotal role in activity-dependent modulation of synaptic structure and neurite development^56,57^. Given that RNMT regulates mRNAs and proteins involved in neurite growth and synaptic function, in phosphorylation-dependent manner, the impact of the RNMT phospho-mutants on neuronal morphology was investigated (Figure 6C). DA neurons expressing GFP-RNMT WT have normal DA neuron morphology, with long neurites. DA neurons expressing GFP-RNMT T317D have reduced Tuj1 expression, which is associated with reduced neurites formation, whereas those expressing GFP-RNMT T317V have bundles of thick, straight and long neurites. Total neurite length (Tuj1 positive) was measured relative to the number of nuclei. Consistent with the increased cytoplasmic neuronal transcripts, DA neurons expressing RNMT T317V have significantly increased neurite growth compared to control DA neurons (Figure 6D).

### RNMT translocation is required to regulate cytoplasmic RNA

To investigate the mechanism by which RNMT regulates cytoplasmic levels of specific RNAs in DA neurons, RNAs regulated in response to RNMT T317D and RNMT T317V were assessed for RNMT binding (RNMT-RNA detection by seCLIP in murine neurons, Figure 7A, Table S1 and S5). DA neurons are human cells and RNMT-RNA interactions were mapped in mouse neurons; only RNAs from genes present in both species were analysed (Figure 7A). For the top translated neuronal mRNAs, there is a positive correlation between RNA enrichment on RNMT and mRNA upregulation in response to RNMT T317V (Figure 7A). There is also a positive correlation between RNA enrichment on RNMT and mRNA down regulation in response to RNMT T317D (Figure 7A). For mRNAs significantly upregulated in response to RNMT T317V, they have enhanced RNMT binding across the transcript compared to unregulated transcripts (Figure 7B). Conversely, mRNAs repressed in response to RNMT T317D have enhanced RNMT binding across the full length of the transcript. Thus, the cytoplasmic mRNAs most responsive to RNMT are those with highest RNMT binding.

**Figure 7:**
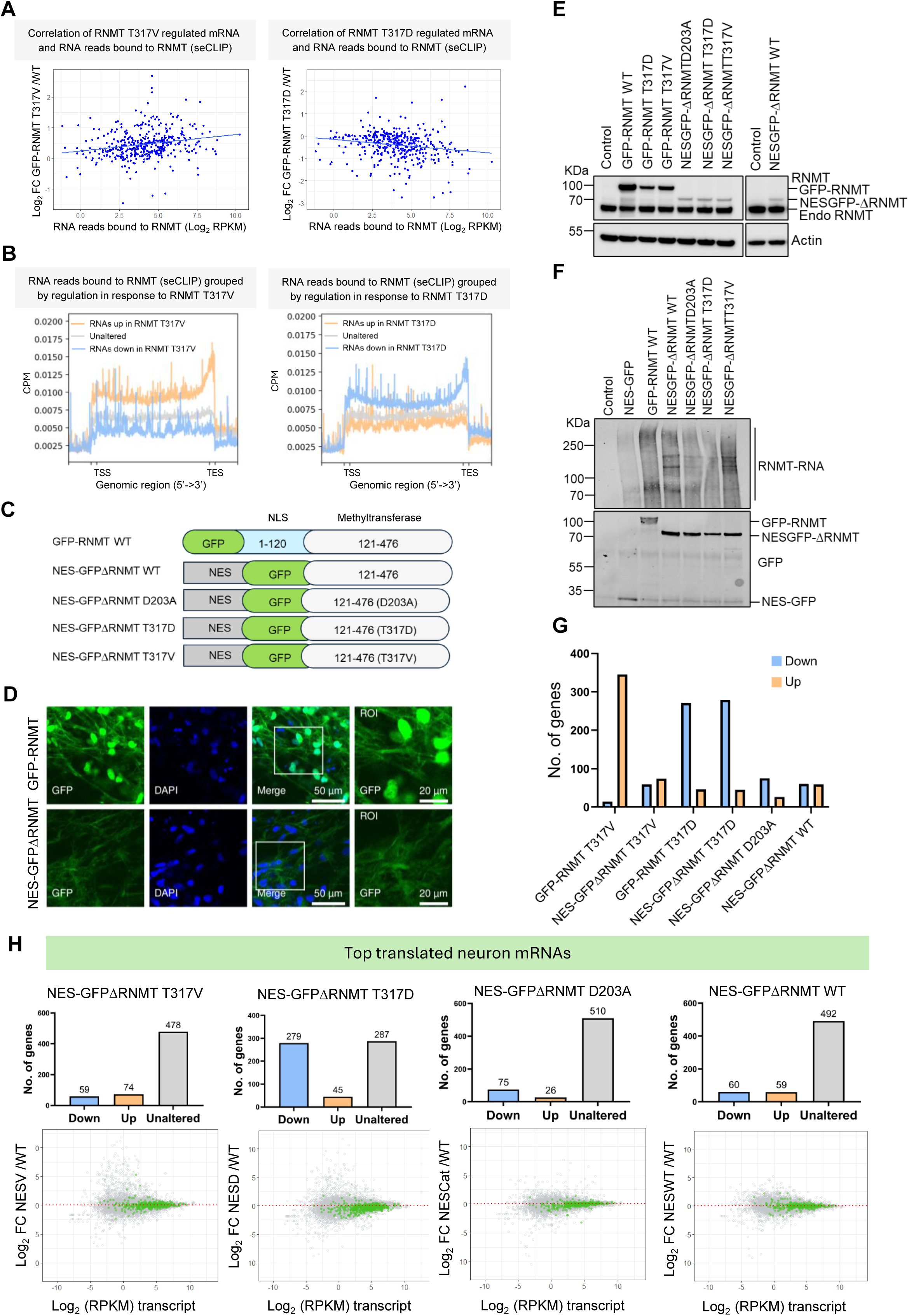
RNMT impact on RNA translocation correlates with RNA binding. A) Correlation of RNMT T317V and T317D-regulated genes and RNMT-RNA interactions. For the top translated neuron mRNAs (Figure 5F), RNA regulation in DA neurons in response to GFP-RNMT T317V or T317D (log_2_fold change) was correlated with RNMT seCLIP reads per gene (RPKM, reads per kilobase per million mapped reads). B) Mapping of RNA which interacts with RNMT for RNMT T317V and T317D regulated genes. For genes significantly upregulated and down regulated in response to RNMT T317V and T317D in DA neurons (Figure 5D), RNMT seCLIP reads were mapped to the genome. C) RNMT mutants used. NES, nuclear export signal. NLS, nuclear localisation signal. GFP, superfolded GFP. D) GFP-RNMT WT and NES-GFP-ΔRNMT WT localisation in hiPSC-derived DA neurons. DAPI nuclear stain (blue). E) Western blot analysis of GFP-RNMT mutants expressed in DA neurons. F) seCLIP analysis of RNMT-RNA interactions in HEK293 cell lines. RNA was UV-cross linked to protein in cells. GFP, GFP-RNMT WT and mutants, were immunoprecipitated via the GFP tag. GFP-RNMT immunoprecipitated from each cell line was balanced by altering the quantity of lysate added to each GFP-IP, see Figure S8. RNA was fragmented, ligated to fluorescent adaptors, resolved by gel electrophoresis and visualised by LI-COR imaging system (upper panel). Immunoprecipitated GFP-RNMT WT and mutants were analysed by western blotting (lower panel). G) For top translated neuron RNAs (Figure 5F, Table S5), number of upregulated and down regulated genes in response to RNMT mutants. H) Scatterplot of transcript log_2_ fold change against transcript abundance in Log_2_ RPKM (reads per kilobase per million mapped reads). Top translated neuron mRNAs highlighted in green ^55^. Bar charts show number of downregulated, unaltered and upregulated transcripts for each cell line.

The features associated with the top RNMT-bound transcripts were examined to probe the gene-specificity of responses. Untranslated regions (UTRs) have been implicated in regulating mRNA stability, localisation, and translation efficiency ^55,58,59^. The top 10% targets identified in RNMT seCLIP in primary neurons (Figure 2A) have longer coding sequence (CDS) and UTRs relative to an equal number of randomly selected seCLIP targets (Figure S10A).

To investigate the impact of RNMT translocation on cytoplasmic RNA level, mutants restricted to the cytoplasm were made. The RNMT N-terminal, non-catalytic domain (1-120) which contains two nuclear localisation signals was replaced by a nuclear export signal (NES), to create the mutant NES-GFPΔRNMT (Figure 7C). NES-GFPΔRNMT was also mutated to either T317V, T317D or D203A (catalytic dead). The NES-GFPΔRNMT proteins were expressed in hiPSCs and differentiated to DA neurons, where they were localised to the cytoplasm (Figure 7D, 7E and S10B-C). RNA interacts with all cytoplasmic-restricted RNMT mutants, confirming that RNMT interacts with RNA in the nucleus and cytoplasm (Figure S10D-E and 7F).

In DA neurons, expression of GFP-RNMT T317V upregulates 345 of the top translated neuronal mRNAs compared to controls (Figure 5F, 7G), whereas the cytoplasmic restricted GFPΔRNMT T317V only upregulates 74 top translated neuronal mRNAs (Figure 7G, H). Thus, nucleus to cytoplasm translocation of RNMT is required for increased levels of neuronal mRNAs in the cytoplasm. Expression of GFP-RNMT T317D results in repression of 271 top translated neuronal mRNAs compared to controls (Figure 5F, 7G). Equivalently, expression of the cytoplasmic-restricted, NES-GFPΔRNMT T317D represses 279 top translated neuronal mRNAs (Figure 7G, H). The phospho-mimic mutation T317D results in RNMT ubiquitination and instability. Since expression of both full length and cytoplasmic-restricted RNMT T317D mutants lead to reduced levels of locally translated neuronal mRNAs, this is consistent with enhanced proteasomal degradation of these proteins leading to active degradation of mRNAs bound to RNMT. In support of this model, the RNAs most enriched on RNMT are the most repressed in response to expression of T317D (Figure 7A and B). Expression of the catalytic dead mutant, NES-GFPΔRNMT D203A, does not result in increased or decreased cytoplasmic mRNAs levels, demonstrating that inhibition of catalysis alone is insufficient to repress RNA (Figure 7G, H).

In summary, here we report that neuron depolarisation results in RNA cap methyltransferase RNMT translocation into the cytoplasm. RNMT binds to RNA in the nucleus and RNMT translocation results in the top translated neuronal mRNAs either being transported or stabilised in the cytoplasm (or both). This mechanism is time-limited by the kinase, CaMKII, which is activated following neuron stimulation. CaMKII-dependent phosphorylation of RNMT results in repression and degradation, and repression of associated mRNAs. By neutralising CAMKII-dependent phosphorylation of RNMT, RNMT-RNA translocation could be observed to be associated with neurite growth control.

## Discussion

Selective control of mRNA dynamics is critical for neuron development and function. A diverse array of RNA binding proteins has been observed to participate in the translocation of specific mRNAs. Here we report that the co-ordinate translocation of the most highly translated mRNAs is directed by redeployment of the RNA cap methyltransferase, RNMT. The master regulator of the response to stimulation, CaMKII, regulates this novel RNMT function. These findings provide insight into the mechanisms that co-ordinate the targeting and regulation of mRNAs at active synapses.

### RNMT function in RNA dynamics in neurons

Neurons are post-mitotic, polarised cells which carry information long distances in the form of waves of depolarisation, followed by the secretion of chemical messengers. Gene expression mechanisms have adapted to the unique function of neurons, translocating mRNAs to their site of requirement in axons and dendrites. Here we report that RNMT is redeployed in neurons to co-ordinate mRNA movement. On stimulation of neurons, RNMT is translocated from the nucleus to the cytoplasm in a calcium-dependent mechanism. Once in the cytoplasm, RNMT binds to actin and other filamentous proteins, localising to punctae in axons and dendrites. Regulated transport of RNMT has not been observed previously and may be a unique feature of neurons and potentially other cells utilising stimulation-dependent calcium signalling.

Translocation of RNMT is associated with increased levels of neuronal mRNAs in the cytoplasm, increased neuronal proteins, including CaMKII, GRM7 and BDNF, and increased neurite growth. Movement of RNMT across the nuclear membrane is specifically required to elevate the abundance of neuronal mRNA in the cytoplasm. Expression of the stabilised mutant, RNMT T317V, which undergoes nuclear to cytoplasmic translocation increases neuronal mRNAs in the cytoplasm whereas expression of NES-ΔRNMT T317V which is retained in the cytoplasm does not. These findings support the model that RNMT predominantly aids mRNA translocation across the nuclear membrane, although RNMT may also contribute to mRNA stability.

Shortly after stimulation, the highly abundant kinase, CaMKII, is activated which co-ordinates cellular responses via phosphorylation of substrates. We demonstrate that RNMT is phosphorylated by CaMKII on T317 which limits RNMT-dependent increases in cytoplasmic mRNAs. RNMT T317 lies adjacent to the active site and its phosphorylation inhibits methyltransferase activity. The phosphomimic RNMT T317D also inhibits catalytic activity, prevents co-factor binding and targets RNMT for degradation. Expression of RNMT T317 T317D in DA neurons reduces levels of locally translated mRNAs, reducing expression of key neuronal proteins and limiting neurite growth. Of note, expression of the catalytic dead RNMT D203A does not repress (or increase) locally translated mRNA levels, indicating that CaMKII-dependent degradation is required to repress mRNA transport, rather than the inhibition of catalytic activity.

### Redeployment of RNMT

RNMT is the earliest evolved RNA cap methyltransferase, present in all eukaryotes. RNMT and homologues complete the ^m7^G RNA cap, which protects RNA and recruits processing and translation factors across eukaryotes^12,13^. In mammalian cells investigated to date, RNMT is predominantly nuclear, with low levels of RNMT observed in the cytoplasm of some cell lines^20^. As with most cellular proteins, RNMT protein is synthesised in the cytoplasm and it has nuclear localisation motifs which direct it to the nucleus ^60,61^. The function of trace cytoplasmic RNMT in cell lines has been ascribed to cytoplasmic RNA cap methylation. In neurons, cytoplasmic RNMT may be involved in recapping, although it is unlikely to be a major function since poly-A mRNA is fully capped^62^. Conversely, RNMT-dependent RNA translocation impacts on the cytoplasmic levels of most of the top 10% of translated neuronal mRNAs.

### CaMKII in stimulation induced gene expression

CaMKII is a major mediator of synaptic plasticity in neurons, including facilitating neuronal mRNAs homeostasis^9,63^. CaMKII is activated on depolarisation and calcium influx, modulating synaptic strength by the direct phosphorylation of AMPA and NMDA receptors. CaMKII has a role in long-term potentiation (LTP) and long-term depression (LTD), including triggering protein degradation by recruiting the proteasome to the synapse^9^. However, with the exception of activating the cytoplasmic polyadenylation element binding protein (CPEB)^43,64,65^, little is known about how CaMKII co-ordinates localised translation. By phosphorylating and inhibiting RNMT following neuron stimulation, CaMKII co-ordinately represses the most highly translated mRNAs encoding neuronal function proteins. This mechanism provides an “off switch” for RNMT function in the cytoplasm. Such off switches are important in neurons for neuronal plasticity^66–68^.

### Stimulation-dependent RNMT export

The stimulation-dependent export of RNMT that we present is the first observation of regulation of RNMT movement. The mechanism of RNMT export is unclear, although RNMT diffuses freely in the nucleus and may simply be released through nuclear pores, which can release proteins larger than 60kDa ^69^. In our analysis, we did not observe CRM1 or other export proteins involved in the TREX pathway to interact with RNMT^70^. Of note, nuclear export of the RNA binding proteins FUS and TDP-43 is shown to be CRM1-independent and likely occurs via passive diffusion^71^. Once released from the nucleus, we observe that RNMT binds to actin and other filament proteins, providing a mechanism for their retention in the cytoplasm. In cortical neurons, RNMT has been identified in proximity to cytoplasmic alpha-synuclein, a marker of synapses^72^.

### Gene specificity of RNMT-RNA interactions

The interaction of RNMT with selective mRNAs in neurons correlates with impact on RNA levels. The transcripts most enriched on RNMT are the most upregulated in the cytoplasm in response to the stabilised RNMT mutant, T317V, and are the most degraded in response to the unstable RNMT phospho-mimic mutant, T317D. Certain RNA binding proteins (RBPs) present in neurons, such as TDP-43, exhibit sequence specificity^73^, whereas others like FUS bind to a range of RNA sequences and structures^74^. RNMT interacts transiently with almost all mRNAs at the 5’ cap during the process of cap methylation and is retained on a large but restricted subset of mRNAs. The mRNAs responsive to RNMT in neurons have a longer 5’ UTR, coding regions and 3’UTRs, than average. Protein-RNA interactions can be governed by nucleotide sequence, nucleotide enrichment and RNA secondary structure. It is likely that a combination of factors, including length of coding region and UTRs, governs which mRNAs interact with RNMT.

In summary, we provide insight into how CaMKII controls proteostasis in neurons, via limiting the function of the RNA cap methyltransferase RNMT following stimulation. We demonstrate that in neurons RNMT is exported from the nucleus into the cytoplasm on stimulation and is redeployed to support translocation of neuronal mRNAs and neurite growth. CaMKII phosphorylation of RNMT inhibits methyltransferase activity, co-factor binding and stability, providing a homeostatic mechanism to limit over-stimulation of neurons. These mechanisms controlling growth could potentially be used to control differentiation in regenerative medicine and brain cancers.

## CONTACT FOR REAGENT AND RESOURCE SHARING

Further information and requests for reagents may be directed to, and will be fulfilled by the corresponding author, Prof. Victoria H. Cowling.

## STATISTICS AND REPRODUCIBILITY

Immunofluorescence (IF) staining of murine brain tissue was performed using two biological replicates and two independent RNMT antibodies. All IF in hiPS cells, DA and primary neurons were replicated independently at least two times. Two independent experiments were performed for depolarisation treatments in primary neurons. In each experiment, hippocampal and cortical neurons were isolated and treated independently as two different sets. RNMT cytoplasmic and nuclear intensities were quantified from the two independent experiments using 5 different fields per each treatment in each set. Neurite length in DA neurons was quantified from three independent neuronal differentiations. RNMT-IP/MS experiments were performed in three different systems (mouse brain, primary neurons and ES cells). RNMT-CaMKII interaction and phosphorylation experiments were replicated independently at least two times. RNA-Seq and CLIP were performed using three biological replicates. The number of replicates processed for (western blot quantifications and qPCR) are reported in the figure legends. The number of MD simulations performed in this study are given in Figure S7C. Graphs and statistical analyses were performed with GraphPad Prism version 10.0.0 and significance was determined using an unpaired student T-test. Values P < 0.05 were considered significant. All data are depicted as means ± standard deviation.

## DATA AND SOFTWARE AVAILABILITY

Sequencing data generated in this study (hiPSC-DA neurons RNA-Seq) were deposited in GEO under the accession number GSE278655 and RNMT seCLIP in primary murine neurons under the accession number GSE198965. All software and packages described in this paper were used according to standard protocols. Custom codes and analysis pipelines were not generated in this paper.

## SUPPLEMENTAL INFORMATION

**Table S1. RNMT seCLIP counts in primary neurons.** RNMT immunoprecipitated in UV-cross-linked primary murine neurons. Raw counts and counts per million (CPM) are given for cross-linked sites for each transcript. The experiment was performed using three biological replicates. For each RNMT IP, only read counts over 10-fold enrichment over IgG control were considered.

**Table S2. RNMT interactome in murine brain.** Proteins identified by mass spectrometry in RNMT immunoprecipitates from brain lysate. For each identified protein, UniProt accession number, coverage, peptide-spectrum match (PSM), and number of unique peptides are given. The Sequest HT score is given. The experiment was performed in two technical replicates. The data were subjected to a 0.05 False Discovery Rate (FDR) at the protein level & peptide spectrum match (PSM) level. Target proteins were filtered using a set of filters in the Proteome Discoverer software (ThermoFisher Scientific).

**Table S3. Cytoplasmic RNMT interactome in primary murine neurons** Proteins identified by mass spectrometry in RNMT immunoprecipitates from control or KCl-treated primary murine neurons. The MaxQuant LFQ (Label-Free Quantification) intensities are given for isotype (IgG) control and RNMT IP samples in each condition. The data were subjected to a 1% False Discovery Rate (FDR) at the protein level & peptide spectrum match (PSM) level.

**Table S4. Summary of RNMT phosphorylation sites identified by MS analysis.** RNMT peptides and phospho-peptides identified by mass spectrometry analysis are given for experiments conducted in this study. The tables include peptide sequences, detected modifications, peptide-spectrum match (PSM), and score.

**Table S5. Summary of cytoplasmic differential gene expression analysis in DA neurons.** RNA_Seq analysis comparing cytoplasmic RNAs from hiPSC-DA neurons control (non-expressing cells) and neurons expressing RNMT mutants. Each row represents a gene. Columns include Gene ID, name and Log_2_ fold change between control cells vs each cells expressing RNMT mutant. Ribo_seq transcripts (neuron translatome mRNA) identified in neuropil (npl) and somata (smt) in (Glock et al. 2021) ^55^ and the top 10% translated mRNA dataset are integrated in the data table. The (Glock et al. 2021) ribo_seq data is also given in a separate sheet. List of genes used in the GO terms analysis are given in a separate sheet. The RNA-Seq was performed using three biological replicates. The differentially expressed transcripts were identified with false discovery rate (FDR)<0.05.

## ACKNOWLEDGEMENTS

We thank Miratul Muqit, Odetta Antico, Sheriar Hormuzdi and Tasha Hunter for their advice and help with primary neuronal cultures. We thank Iain Porter and Jason Swedlow for providing reagents and support with the hiPSC-DA neuron cultures. We thank Don Tennant, Sarah Thomoson and Claire Johnstone for their help with resources. We thank Catherine Winchester, Peter Thomason, Claire Mitchell, and Beatrice Bottura (CRUK Scotland Institute) for helpful comments on the manuscript. We thank the microscopy facilities at the University of Dundee School of Life Sciences (SLS) and CRUK Scotland Institute. We thank Dundee FingerPrints proteomics facility for help with the proteomics processing and analysis. We thank Tayside Centre for Genomic Analysis facility for the seCLIP sequencing. We thank the staff at University of Dundee, CRUK Scotland Institute and the Cowling lab for their help and support.

## AUTHOR CONTRIBUTIONS

Conceptualization RA, VHC; Methodology all; Formal Analysis RA, SL, MB, LH, LD, FH, AP; Investigation RA, SL, MB, LH, FH; Writing, Review & Editing all; Visualization all, Funding Acquisition VHC, AVP, AIL,

---

**Figure S1.**
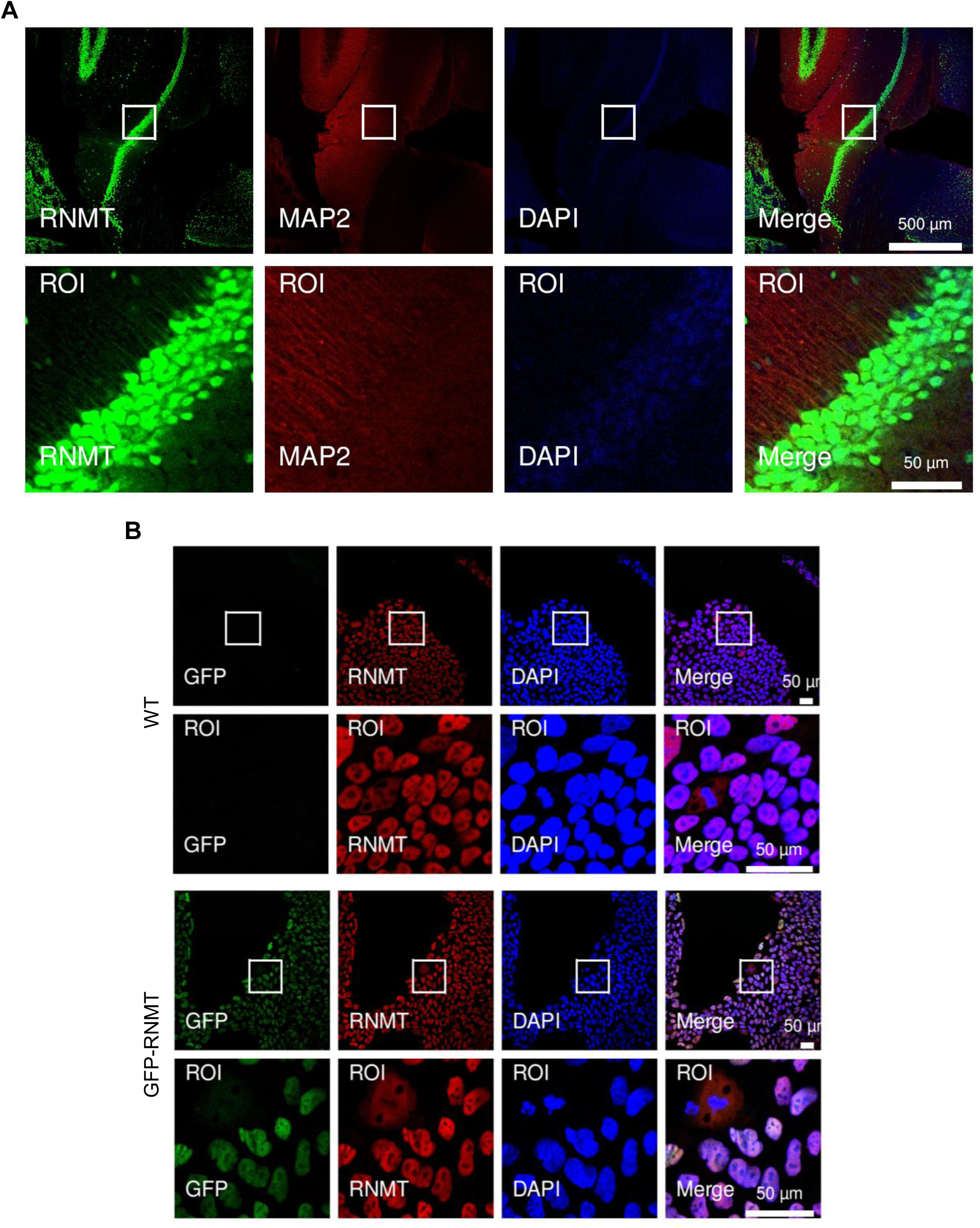
RNMT localisation in neurons (related to Figure 1) A) Immunoflourecence (IF) staining in adult mouse brain sections using polyclonal anti-mouse RNMT raised in rabbit. RNMT (Green), DAPI (blue), MAP2 (red). (B) RNMT IF using polyclonal anti-human RNMT raised in sheep and GFP-RNMT localisation in undifferentiated hiPSC, GFP (green), RNMT (red), DAPI (blue).

**Figure S2.**
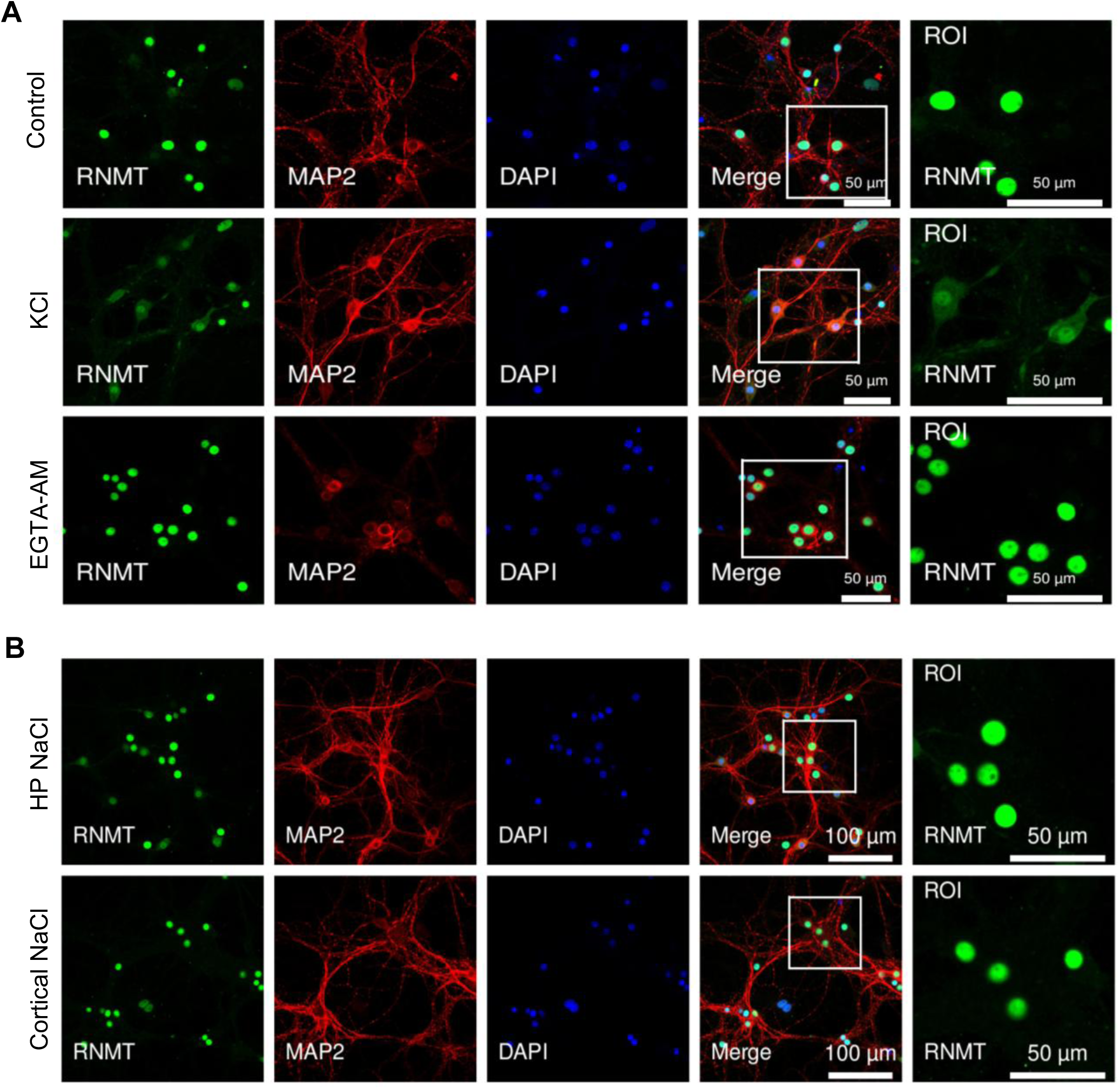
Depolarisation of cortical neurons induces RNMT translocation from the nucleus to the cytoplasm (related to Figure 1) A) RNMT detected by IF in primary cortical neurons (DIV16) following incubation with 50 mM KCl for 5 mins followed by 30 mins recovery or 50 μm EGTA-AM incubation for 30 mins. RNMT (green), MAP2 (red), DAPI (blue). See Figure 1B for RNMT IF in primary hippocampal neurons. B) RNMT detected by IF in primary hippocampal (HP) and cortical neurons (DIV16) following 50 mM NaCl for 5 mins, followed by 30 mins recovery. RNMT (green), MAP2 (red), and DAPI (blue).

**Figure S3.**
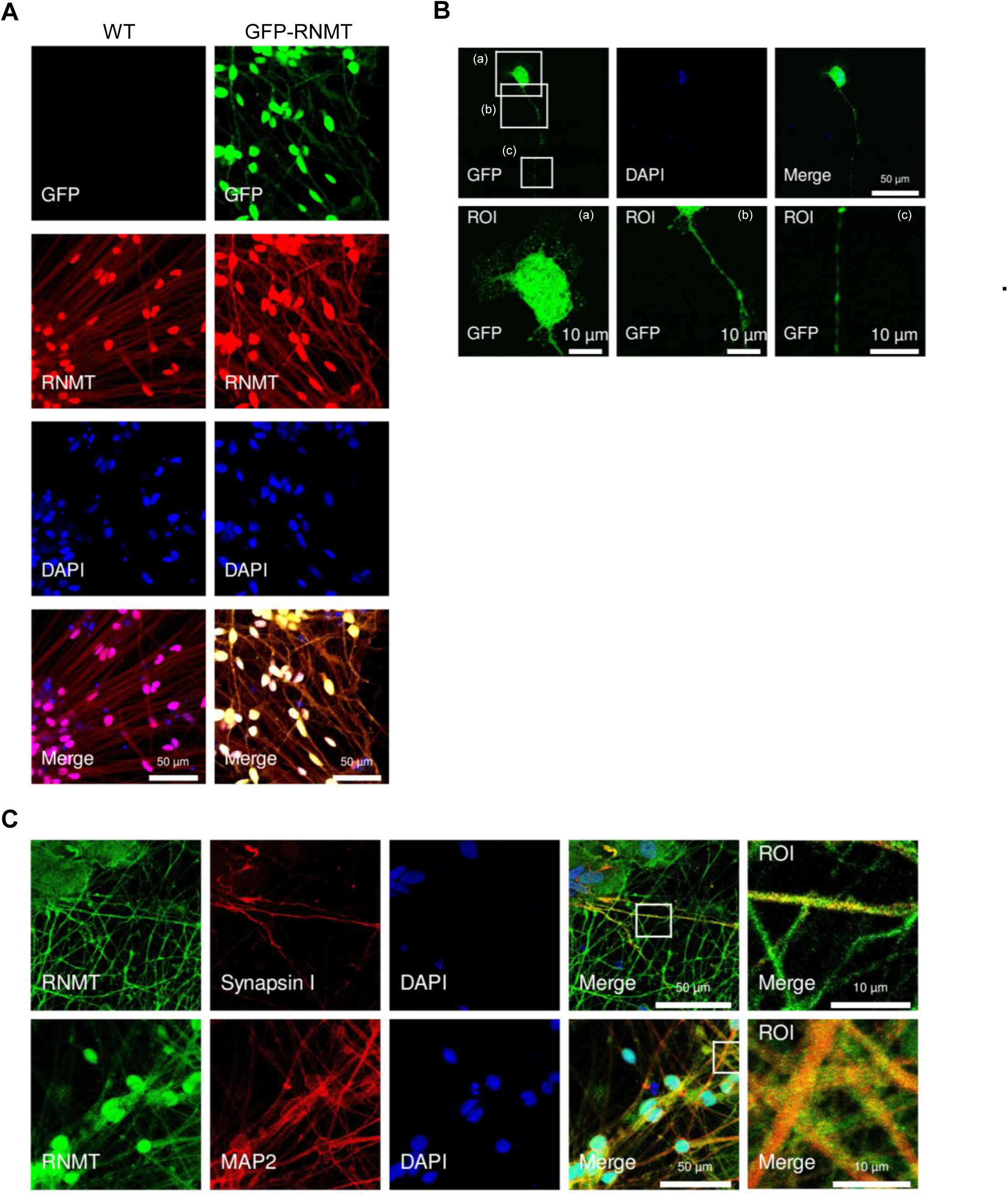
RNMT localises to neurites and dendrites (related to Figure 1) (A) Endogenous RNMT and GFP-RNMT localisation in hiPSC-derived Dopaminergic (DA) neurons. GFP (green), RNMT staining using polyclonal anti-human RNMT raised in sheep (red), DAPI (blue). B) Primary murine neurons transfected with GFP-RNMT visualised at 7 days in vitro (DIV). GFP-RNMT expression visualised directly by fluorescence microscopy. C) RNMT in Synapsin I-positive neurites and MAP2-positive dendrites in DA neurons. RNMT staining using polyclonal anti-human RNMT raised in sheep (green), MAP2 (red), Synapsin I (red), DAPI (blue).

**Figure S4.**
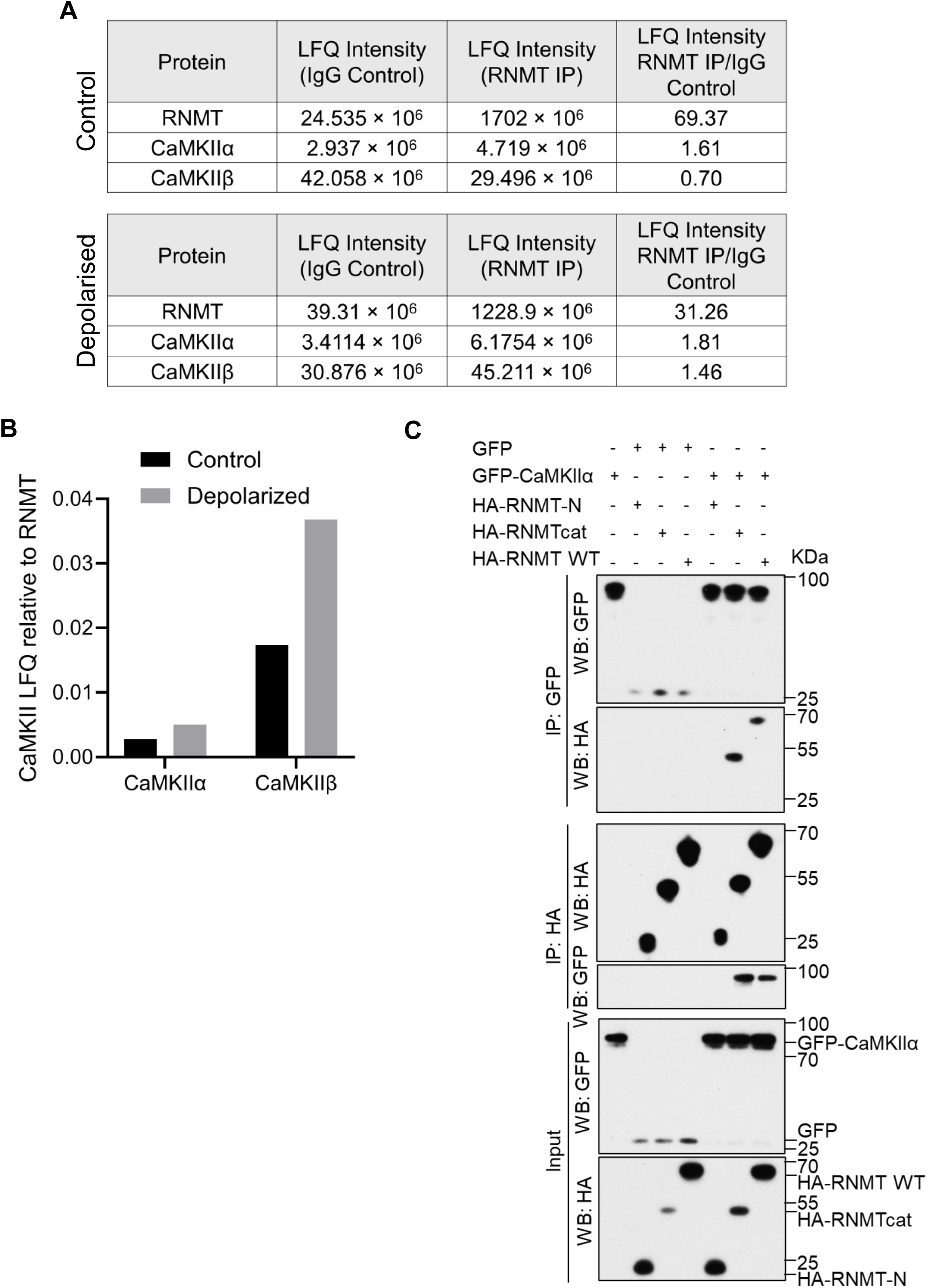
RNMT-CaMKII interaction in primary neurons (Related to Figure 2) A) RNMT was immunoprecipitated (IP) from the cytoplasmic fraction of untreated control or KCl-depolarised primary cortical neurons. Mass spectrometry analysis was performed on RNMT IPs. Maxquant LFQ intensity for RNMT and CaMKII proteins reported. B) For CAMKIIα and CAMKIIβ, LFQ intensity normalised to RNMT LFQ intensity. C) HA-RNMT, HA-RNMT-N (1-120) and HA-RNMTcat (121-476) were co-expressed with GFP or GFP-CaMKIIα in HEK293 cells. HA-RNMT and GFP-CaMKIIα were immunoprecipitated via their tags. Western blot analysis was performed.

**Figure S5.**
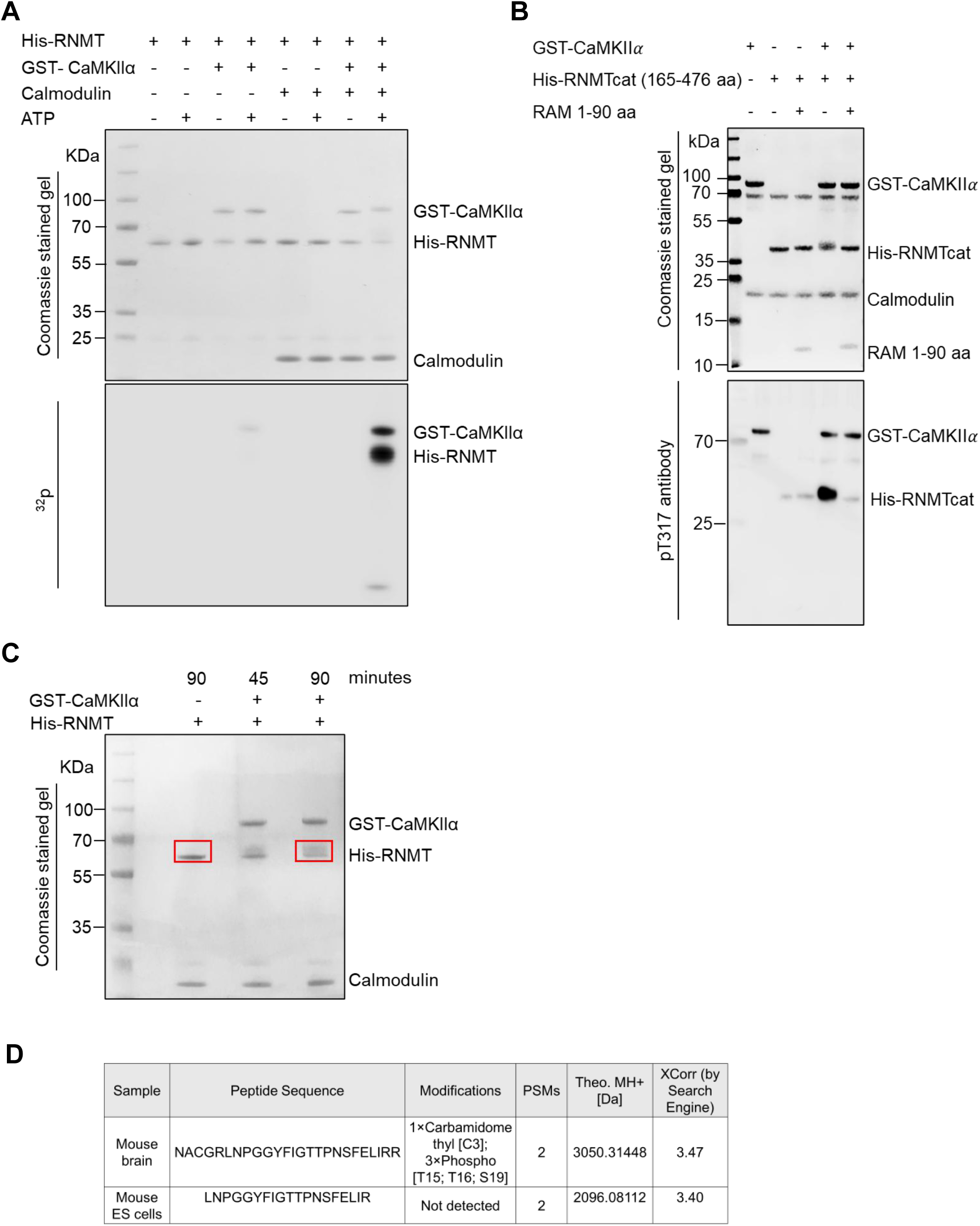
RNMT phosphorylation on T317 (Related to Figure 3) A) Recombinant full length His-RNMT was incubated with activated GST-CaMKIIα, calmodulin, and ^32^P-γATP for 90 min. Samples were resolved by SDS-PAGE. Labelled bands were visualised by phospho-imaging. Recombinant proteins were visualised by Coomassie Blue stain. B) Recombinant RNMT and RNMT-RAM1-90 were incubated with active GST-CaMKIIα and ^32^P-γATP for 90 minutes. RNMT pT317 was visualised by western blotting with anti-RNMT pT317 antibody. C) Coomassie Blue-stained gel of CaMKII-phosphorylated His-RNMT. Indicated bands were excised and analysed by MS for modification site detection. D) RNMT phosphorylation sites analysed by MS in immunoprecipitated RNMT from mouse brain and embryonic stem cell (ESC) lysate. pT317 is detected in brain lysate but not in ESC.

**Figure S6.**
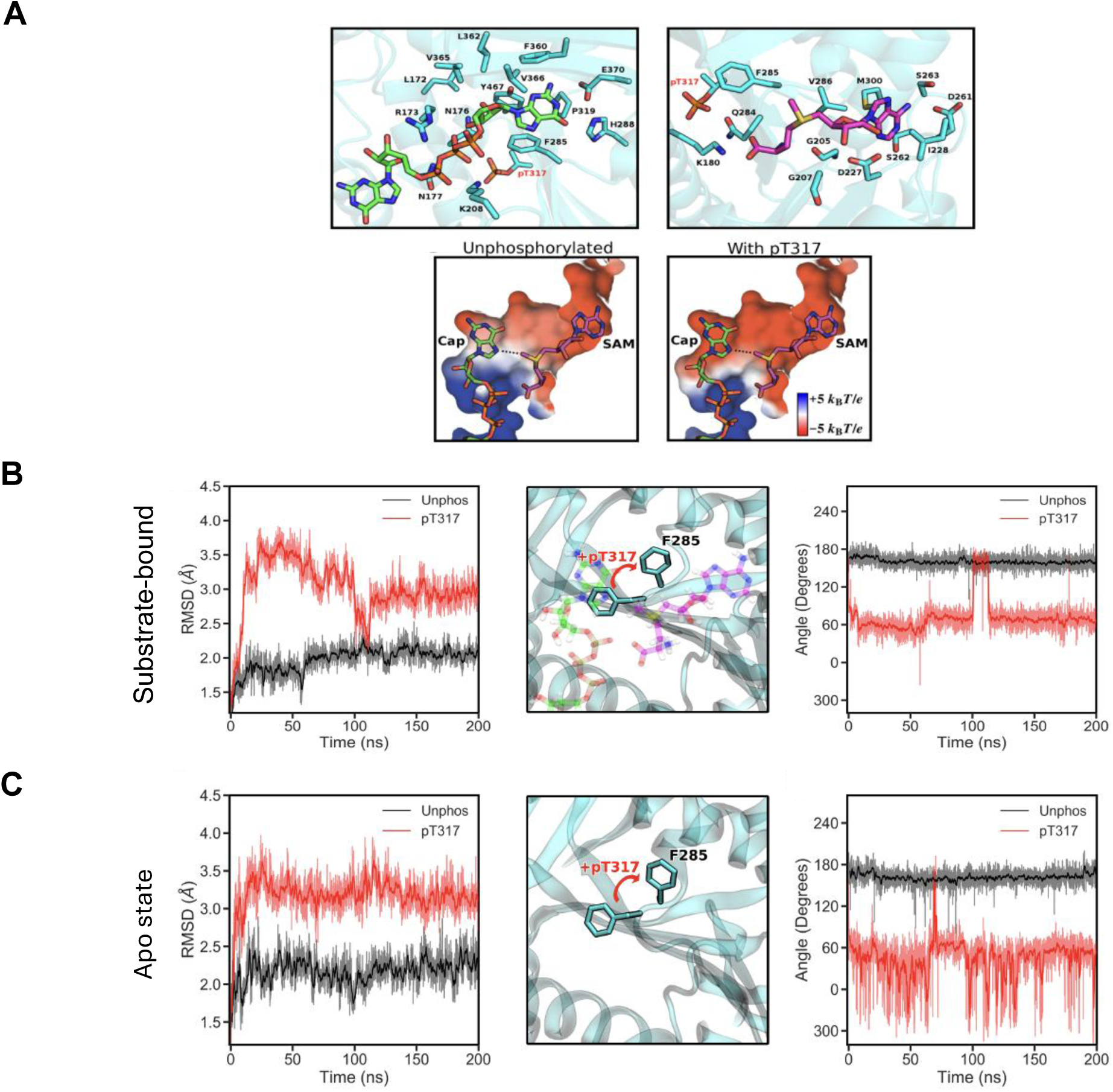
The impact of T317 phosphorylation on the cap-binding pocket and simulations of T317 mutations (Related to Figure 4) A) RNMT substrate binding sites in relation to pT317. The coordination of the cap (top left), and SAM, (top right), as described by Bueren-Calabuig et al.^44^ Residues that coordinate the substrates are indicated in black. T317 phosphorylation is modelled and labelled in red. The electrostatic potential surface representation of the unoccupied substrate binding sites, with RNMT modelled in the unphosphorylated and T317 phosphorylated states (bottom). pT317 is modelled in the -2 state. The substrates are superimposed into the binding pockets as a reference. (B-C) The effect of T317 phosphorylation on the cap-binding pocket in (B) the substrate-bound or (C) apo state. Time-evolution RMSDs of the core residues that coordinate the cap (G0) in the RNMT active site in the substrate-bound states (see Figure S7C, Systems 1 and 2) (top) and apo state (see Figure S7C, systems 3 and 4) (bottom) is shown in the left panel. Residues included in the selection are taken from Panel A, excluding K208, V365 and V366 due to their high flexibility. Representative snapshots of the RNMT active site showing the conformational change of F285 upon T317 phosphorylation are given in the middle panel. Snapshots of the systems were taken at 150 ns. The time-evolution of the dihedral angle that describes the rotation of the F285 sidechain (C-Cα-Cβ-Cγ) is shown in the right panel.

**Figure S7.**
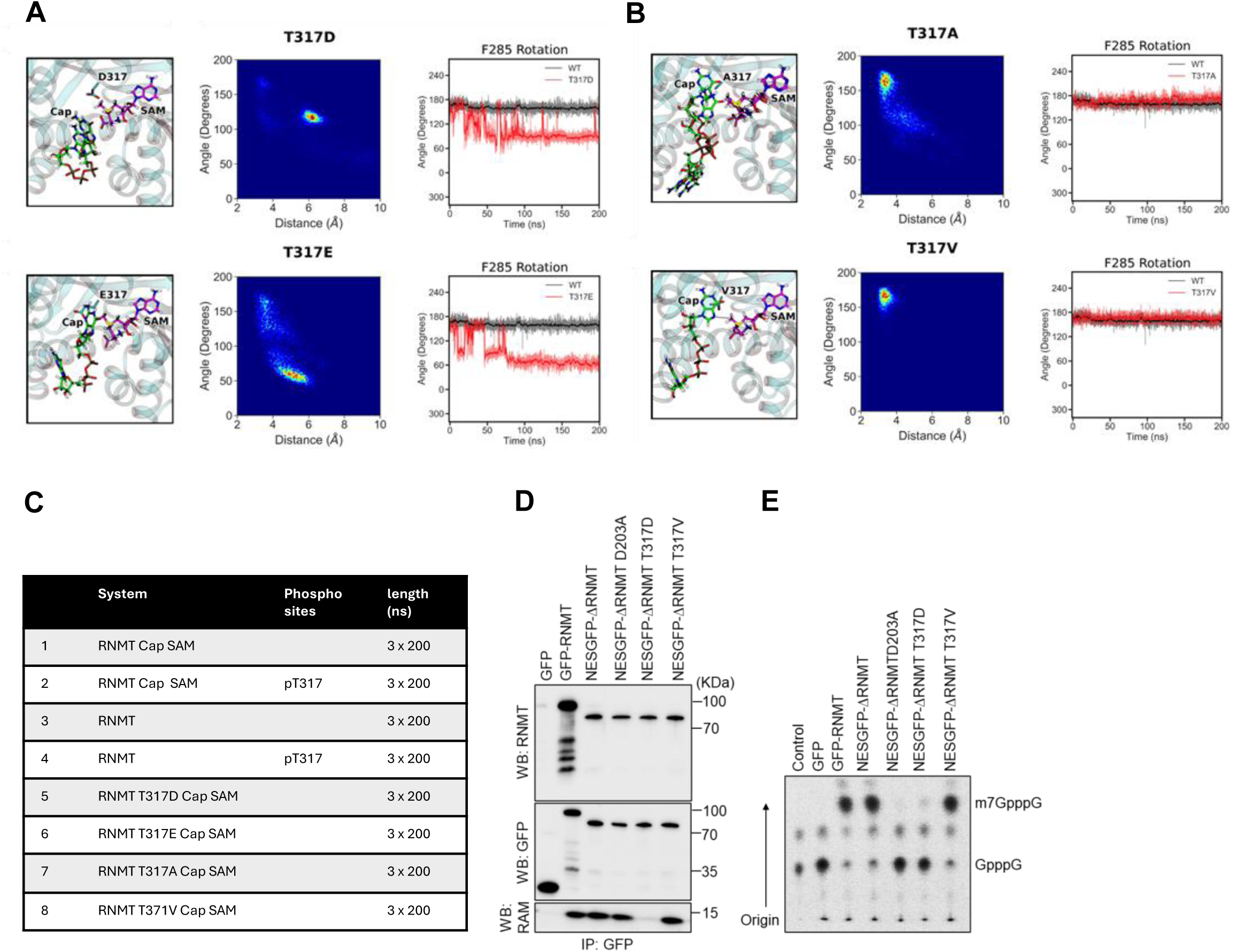
(Related to Figure 4) Simulations of T317 mutations. A) Molecular dynamics simulations. Substrate coordination for the two T317 phospho-mimetic mutants (T317D (top) and E (bottom)). Representative snapshots showing the most occupied conformation of the RNMT substrates during MD simulations are shown in the left panel. Heatmaps displaying the distance (cap G0 N7 – SAM methyl group) and angle (cap G0 N9 – G0 N7 – SAM methyl group) of the RNMT reactive groups are given in the middle panel. The time-evolution of the dihedral angle that describes the rotation of the F285 sidechain (C-Cα-Cβ-Cγ) is shown in the right panel. For both systems, the cap becomes destabilised, causing a deviation in the optimal coordination required for catalysis. The rotation of F285, observed in the destabilisation of the cap pocket during T317 phosphorylation, is also observed in both phospho-mimetic mutant systems. B) Substrate coordination for the two T317 phospho-dead mutants (T317A (top) and V (bottom)). Data panel as in (A). In the T317A simulation, although the substrates remained predominantly in the active conformation for the duration of the simulations, there was some destabilisation of the active site. In particular, the cap occupied inactive conformations during the simulations. Observing the destabilisation of the active site on the relatively short timescale of these simulations suggests that the basal activity of RNMT might be significantly affected by the T317A substitution. In contrast to the T317A mutant, the T317V simulations displayed a stable active site that is comparable to the wild-type RNMT system. In all repeats, the substrates remained in the active conformation and there was no rotation of F285. C) Summary of the molecular dynamics simulations performed for RNMT T317 simulations. D) RNMT mutants transiently expressed in HEK293 were immunoprecipitated and analysed by western blot. E) RNA cap methyltransferase activity in immunoprecitates was determined by *in vitro* assay. Resultant m7GpppG (methylated cap) and GpppG (unmethylated substrate) were separated by thin layer chromatography and detected by phospho-imaging.

**Figure S8.**
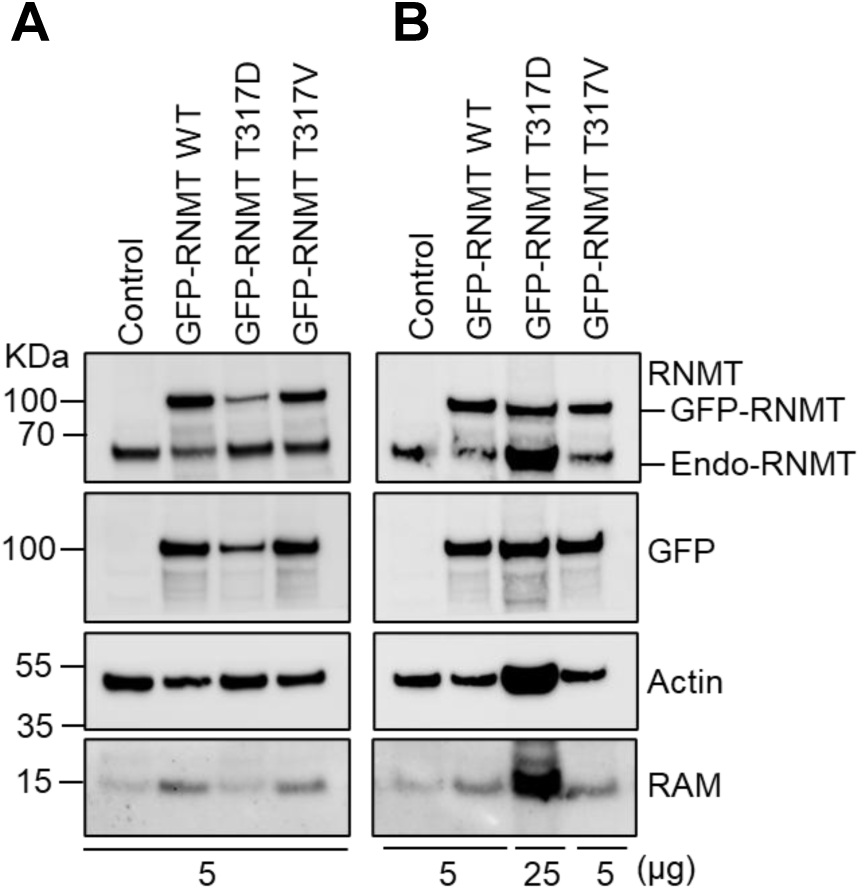
GFP-RNMT WT and mutants expression in HEK293 cells (related to Figure 4) HEK293 cell lines were generated to express GFP-RNMT WT, T317D and T317V. (A) 5 μg cell extracts was analysed by western blotting. GFP-RNMT T317D was expressed at 5-fold less than GFP-RNMT WT and T317V. (B) 5 μg HEK293 cells expressing GFP-RNMT WT and T317V and 25 μg cell extracts expressing T317D analysed by western blotting.

**Figure S9.**
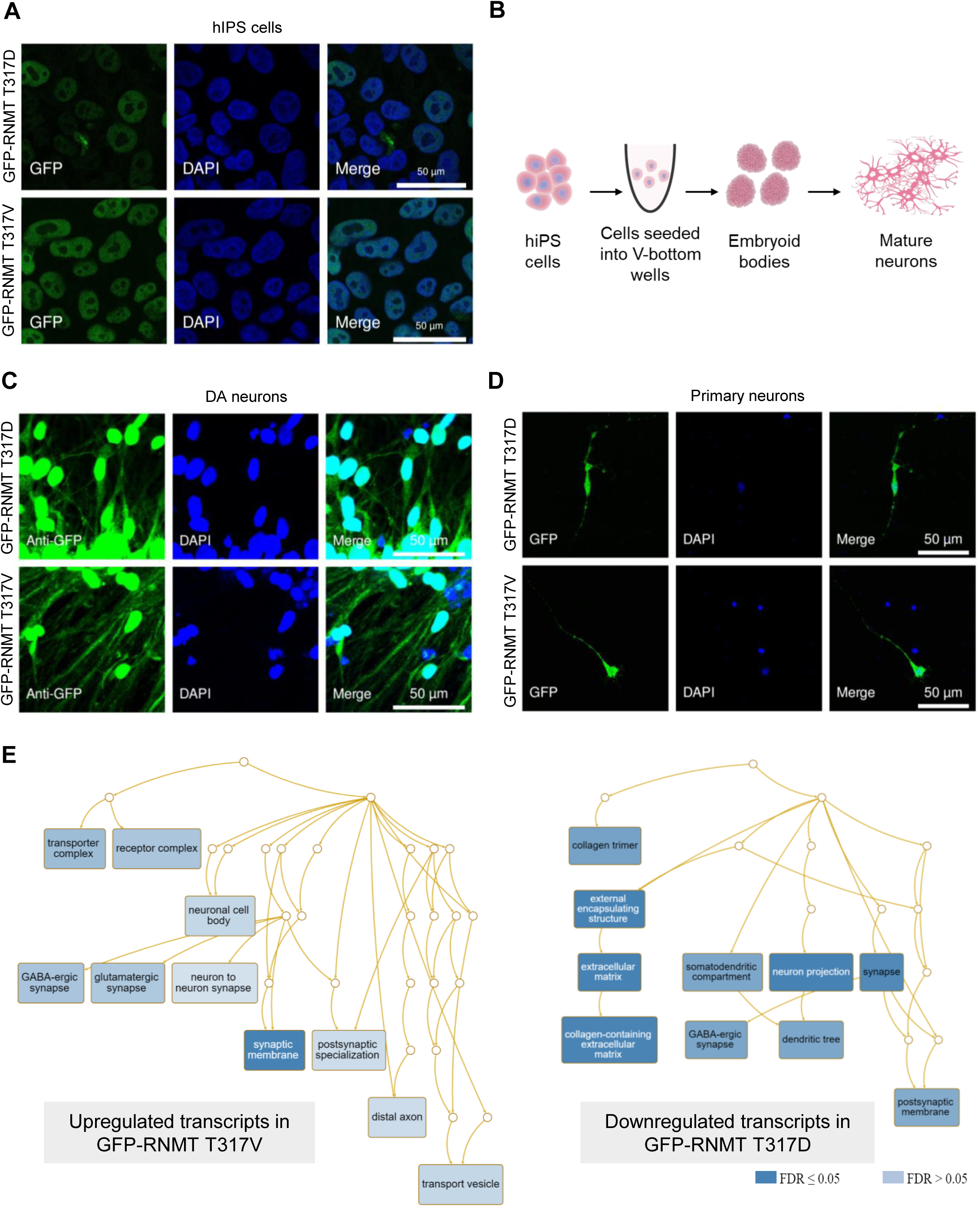
Establishing RNMT T317 phospho-mutants (Related to Figure 5) A) GFP-RNMT T317D and T317V localisation in undifferentiated hiPSC B) Representation of hiPSC differentiation into DA neurons. Embryoid bodies (EBs) generated by seeding hiPS cells in V-bottom well plates were used for differentiation. C-D) GFP-RNMT T317D and T317 localisation in C) hiPSC-derived DA neurons and D) primary hippocampal neurons (DIV 7). GFP-RNMT (green) and DAPI (blue). E) GO terms analysis of the top 20% downregulated transcripts in GFP-RNMT T317D-DA neurons (left panel) and top 20% upregulated targets in GFP-RNMT T317V-DA neurons (right panel) compared to wild-type cells.

**Figure S10.**
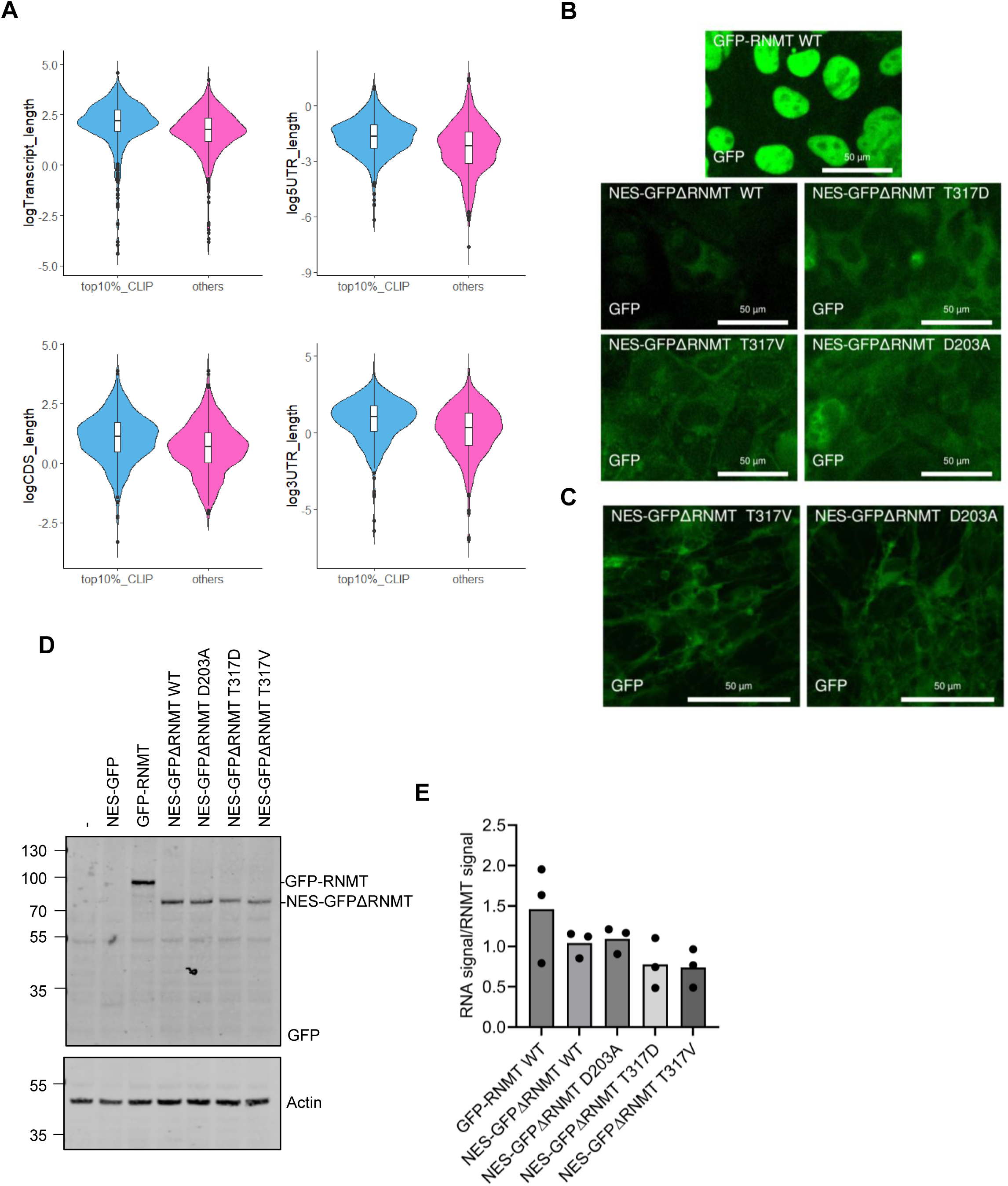
Cytoplasmic RNMT T317 phospho-mutants (Related to Figure 7) A) Features of top RNMT-bound transcripts in primary neuron. Top 10% targets (683 genes) identified in RNMT seCLIP analysis (see Figure 2A) were compared to an equal number of randomly selected seCLIP targets. Violin plots show the distribution of the transcripts, UTRs and coding sequence (CDS) length. B) GFP-RNMT WT and NES-GFPΔRNMT mutants expressed in hiPSC. GFP, green fluorescence. C) NES-GFP-ΔRNMT T317V and D203A localisation in DA neurons derived from hiPSC. GFP signal was visualised directly by fluorescence microscopy. D) GFP-RNMT and NES-GFPΔRNMT mutants expressed in HEK293 cells. E) GFP-seCLIP analysis was performed on GFP-RNMT and NES-GFPΔRNMT mutants expressed in HEK293 cells (see Figure 7F). RNA immunoprecipitated with GFP-RNMT mutants in HEK293 cells is quantified for 3 independent experiments.

## STAR+METHODS

## KEY RESOURCES TABLE

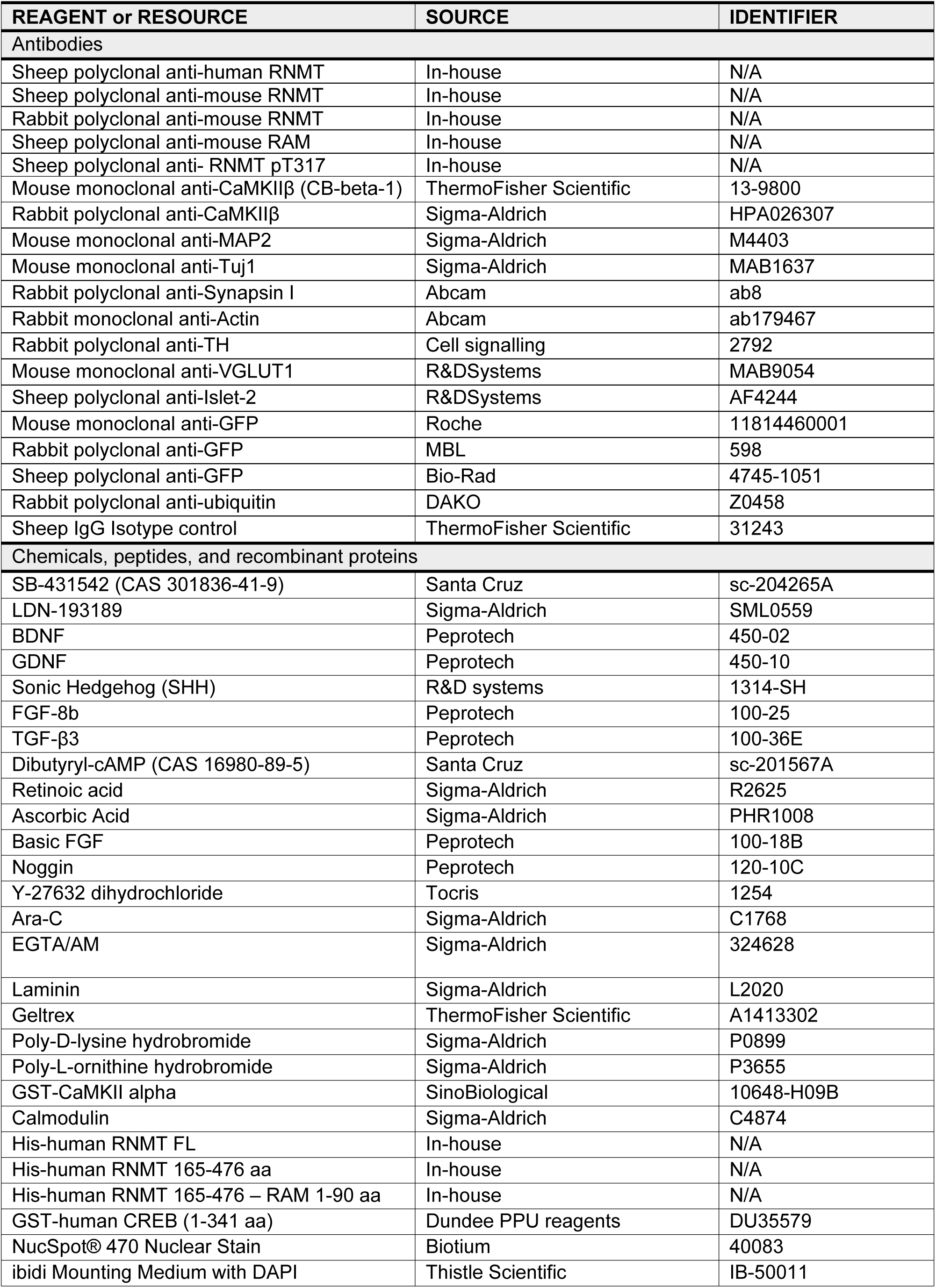
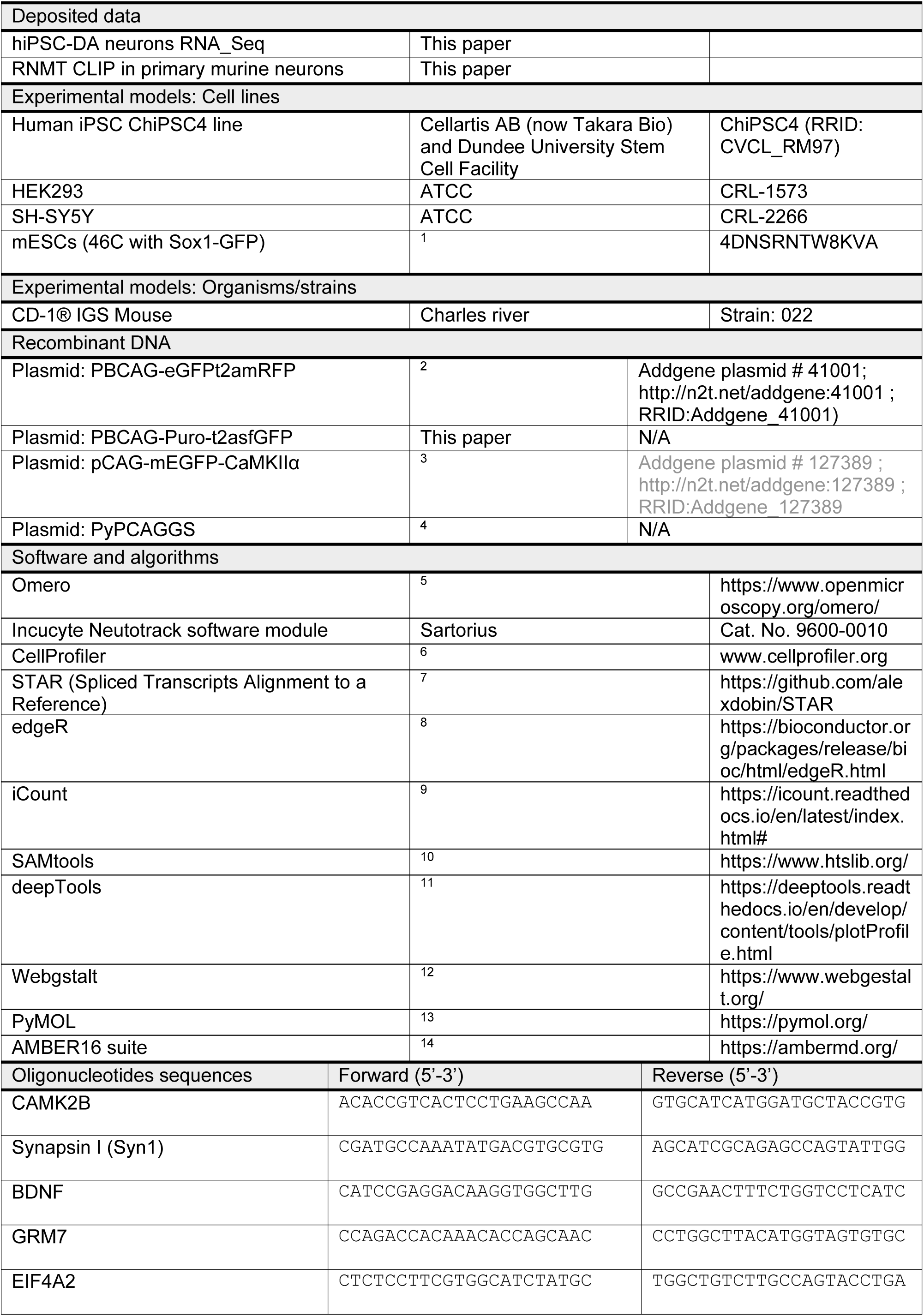
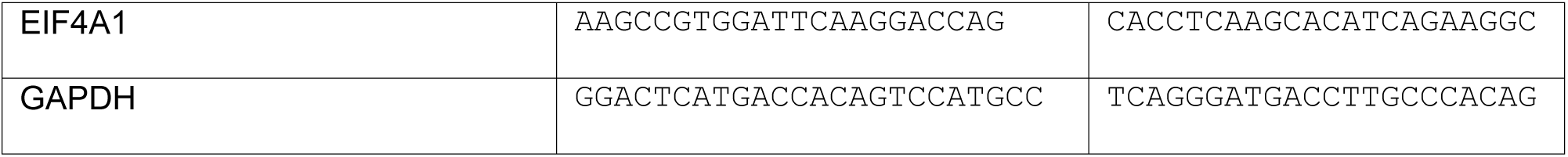

## METHOD DETAILS

### Cell Culture

The human iPSC line ChiPSC4 (Takara Bio) was used to generate RNMT mutant cell lines. ChiPSC4 culture and maintenance was performed according to ^15^ with slight modifications. Briefly, cells were cultured on Geltrex matrix (ThermoFisher Scientific) coated plates (20 μg/ cm^2^) with in-house prepared TeSR medium supplemented with bFGF (30 ng/ml) and human Noggin (10 ng/ml, Peprotech)^16^. For passaging, the medium was supplemented with Rho kinase inhibitor Y-27632 (10 μM, Tocris). HEK293 and SH-SY5Y cells were cultured in DMEM supplemented with 10% fetal bovine serum, 2 mM L-glutamine, 100 U/ml penicillin, and 100 mg/ml streptomycin. Plasmid transfections in HEK293 and SH-SY5Y cells were performed using Lipofectamine 2000 (ThermoFisher Scientific). Mouse embryonic stem cells (mESCs) were cultured as described in ^4^. Briefly, ES cells (expressing Sox1-GFP) were cultured on 0.1% gelatin-coated dishes in Glasgow minimal essential medium (GMEM) (Sigma), 10% knockout serum replacement, 1% modified Eagle’s medium (MEM) non-essential amino acids, 1 mM sodium pyruvate (ThermoFisher Scientific), 0.1 mM 2-mercaptoethanol, and 100 U/ml recombinant human leukaemia inhibitory factor (LIF). All cell lines were cultured at 37°C in the presence of 5% CO2.

### Primary neuronal culture

E17-18.5 mice embryos (Crl:CD1(ICR), Charles River Laboratories) were used to isolate primary cortical and hippocampal neurons. Dissected cortices and hippocampi were dissociated using 1.5 mg/ml Papain in HBSS for 30 minutes at 37°C. Isolated neuronal cells were plated onto 0.1 mg/ml poly-D-lysine-coated dishes and maintained in neurobasal medium supplemented with 1X B27, 2 mM GlutaMAX, 100 U/ml penicillin, and 100 mg/ml streptomycin. Cells were treated with 5 μM Cytosine arabinoside (AraC) on DIV 2 for 24 hours to inhibit proliferating cells. Cells were allowed to grow to form mature neurons and harvested at DIV15-18 or as indicated. DNA transfection of primary neurons (DIV 4-6) was performed using Lipofectamine 2000 (ThermoFisher Scientific). Cultured neurons were maintained at 37°C in the presence of 5% CO2.

### Plasmid subcloning

The GFPt2aRFP cassette in the piggyBac vector (PBCAG-eGFPt2amRFP, Addgene 41001) was replaced with a puro-t2a-XhoI_GFP_BglII cassette to allow for puromycin selection. The sequences of superfolder (sf)GFP-RNMT WT, and mutants were synthesised as DNA fragments by ThermoFisher GeneArt, and were subcloned into the modified piggyBac vector between XhoI and BglII sites. Subcloned sequences were verified by Sanger sequencing. For cytoplasmic restricted constructs, the N terminus of the catalytic domain of RNMT (121-476 aa) was tagged with the Xenopus MAPKK nuclear export signal (32 ALQKKLEELELDE 44) ^17^ fused to sfGFP.

### hiPSC transfection and generation of ChiPSC4-sfGFP-RNMT lines

To make ChiPSC4 sfGFP-RNMT mutant cell lines, 1.5 × 10^6^ cells were transfected with piggyBac (1 μg) and transposase (0.150 μg) (Cambridge Bioscience) constructs using the Neon transfection system (ThermoFisher Scientific). After transfecting cells with either 1100 or 1150 Volts-single pulse for 30 ms, cells were allowed to recover for 24 hours prior to selection for 72 hours with puromycin (1 µg/ml). Emerging clones were passaged and screened for the presence of GFP-RNMT expression using microscopy and western blotting.

### hiPSC neuronal differentiation

hiPSC derived dopaminergic, motor, and forebrain neurons were generated as previously described^18^. Briefly, embryoid bodies (EBs) were generated by spinning down cells (18,000 cells/well) in V-bottom 96-well plates (Corning). EBs were seeded onto poly-L-ornithine (15 μg/ cm^2^; Sigma)/ laminin (1 μg/ cm^2^; Sigma)-coated dishes. Neural induction was initiated by applying a dual SMAD inhibition protocol in the presence of TGF-β inhibitor SB431542 (Santa Cruz) and BMP signalling inhibitor LDN193189 (Sigma) inhibitors to generate neural precursors. Dopaminergic neuronal differentiation was induced using neurobasal medium with 1X B27, 1X N-2, 2 mM GlutaMAX (base medium) supplemented with 20 ng/ml human sonic Hedgehog (SHH; R&DSystems), 100 ng/ml fibroblast growth factor 8b (FGF-8b, PeproTech) 200 μM ascorbic acid (AA, Sigma-Aldrich) and 20 ng/ml brain-derived neurotrophic factor (BDNF, PeproTech) for 10 days. To allow for neuronal growth, the base medium was then supplemented with 200 μM AA, 20 ng/ml BDNF, 20 ng/ml glial cell-derived neurotrophic factor (GDNF, PeproTech), and 0.2 mM cyclic AMP (cAMP, Santa Cruz). Cells were then maintained for another 10 days before harvesting. For motor neuron induction, neurobasal base medium was supplemented with 20 ng/ml BDNF, 200 μM AA, 20 ng/ml SHH and 0.5 μM retinoic acid (RA) for 20 days. Forebrain neurons were induced by maintaining neural precursors in neurobasal base medium, supplemented with 20 ng/ml BDNF and 20 ng/ml GDNF for 20 days. The medium was changed every 2-3 days until cell harvesting.

### Primary neurons depolarisation

Primary neurons were activated for 5 minutes by adding 50 mM KCl in fresh medium. Subsequently, the medium was replaced with conditioned medium, and cells were allowed to recover for 30 minutes before harvesting. For Ca2+ chelating treatments, 50 μM EGTA-AM (Sigma-Aldrich) or an equal volume of vehicle (DMSO) was incubated with cultures for 30 minutes before harvesting.

### Protein extraction and western blotting

Cells and brain tissue were lysed in 10 mM Tris (pH 7.05), 50 mM NaCl, 50 mM NaF, 10% glycerol, 0.5% Triton X-100 supplemented with 1 mM DTT, 10 mM leupeptin, 1 mM pepstatin, 0.1 mg/mL aprotinin, and phosphatase inhibitor PI2 and PI3 cocktail (Sigma). For western blot analysis, lysates were boiled in Laemmli buffer at 95°C for 10 minutes. Equal amount of total protein was loaded onto an SDS-PAGE gel and transferred to activated PVDF membrane using wet transfer system. Blot detection was carried out using horseradish peroxidase (HRP) conjugated secondary antibodies and enhanced chemiluminescence (ECL) (ThermoFisher Pierce). Bands quantification was carried out using Image Lab software (Bio-Rad). Actin or GAPDH were used as loading control to check for sample integrity.

For mass spectrometry analysis of RNMT interactions in mouse neuronal tissues and ESCs, brain tissue was lysed in 20 mM HEPES pH 7.4, 50 mM NaCl, 0.5% Triton, 30 mM Na4 pyrophosphate, 50 mM NaF, 10% glycerol, 5 μM ZnCl2, 0.5 mM DTT supplemented with 10 mM leupeptin, 1 mM pepstatin, 0.1 mg/mL aprotinin, and phosphatase inhibitor PI2 and PI3 cocktail (Sigma). To generate cytoplasmic extracts in primary neurons, cells were depolarised as previously described. Subcellular fractionation was carried out using a modified protocol from^19^. Briefly, cells were collected by centrifugation and resultant pellets resuspended in 250 μl HLB (10 mM Tris pH 7.5, 10 mM NaCl, 3 mM MgCl2, 0.5% (v/v) NP40, 10% (v/v) Glycerol, 0.32 M sucrose) containing 10% RNase inhibitors (RNasin Plus, ThermoFisher Scientific). Samples were incubated for 10 minutes on ice and then spun at 1000g for 3 minutes at 4°C. The supernatant was layered over 250 μl of a 1.6 M sucrose cushion and spun at 21,000g for 5 minutes. For Sox1-GFP mESCs subcellular fractionation, cells were lysed in Buffer A (10 mM HEPES pH7.9, 10 mM KCl, 1.5 mM MgCl_2_). Pelleted nuclei were then re-suspended in Buffer C (20 mM HEPES pH 7.9, 420 mM NaCl and 1.5 mM MgCl_2_) supplemented with 1000U/ml benzonase, protease and phosphatase inhibitors. The lysate was then incubated for 10 minutes at 4°C and another 10 minutes at 37°C before spinning down at 4000g to collect the nuclear supernatant. 0.5% Triton was added to the supernatant before proceeding with RNMT immunoprecipitation.

### Immunoprecipitation (IP)

Immunoprecipitations of mouse brain homogenate were performed using 1 μg of sheep anti-RNMT (in-house) or IgG control (ThermoFisher Scientific) added to 3 mg of tissue lysate with 20 μl of protein G Dynabeads (ThermoFisher Scientific). After 2 hours incubation at 4°C, beads were washed with lysis buffer and boiled in 2X Laemmli buffer for 10 min. For HA and GFP immunoprecipitations in HEK293 and SH-SY5Y cells, 20 μl anti-HA magnetic beads (ThermoFisher Pierce) or 1 μg of anti-GFP (Roche) with 20 μl of protein G Dynabeads (ThermoFisher Scientific) were added to 1 mg of cell lysate. After an overnight incubation at 4°C, beads were washed with lysis buffer and eluted by boiling in 2X Laemmli buffer. For MS-IPs in brain tissue samples, the lysate was pre-cleared with protein G Sepharose beads for 1 hour at 4°C before proceeding with the IPs as described above. To detect RNMT pT317 by western blots, 2 μg of sheep anti-RNMT (in-house) was added to 5 mg brain lysate with 30 μl of protein G Dynabeads (ThermoFisher Scientific). For MS-IPs in mESC, 3 μg of anti-RNMT was added to 3 mg of cell lysate with 20 μl protein G Dynabeads for each IP. Samples were incubated overnight at 4°C then beads were washed with lysis Buffer C supplemented with 0.5% triton before eluting by boiling in 2X Laemmli buffer. For MS-IPs in primary neurons, lysates were pre-cleared with 10 μl of protein G Dynabeads. 5 μg of anti-RNMT or IgG control was added to 1 mg lysate with 50 μl protein G Dynabeads (ThermoFisher Scientific) for each IP. After 3 hours incubation at 4°C, beads were washed in fresh HLB with 50 μM salt (no sucrose) and eluted by boiling in 20 μl 1X Laemmli buffer.

### Mass spectrometry

Lysates and IPs were prepared as previously described. Endogenous RNMT was immunopurified from brain and mouse ES cells with an anti-RNMT antibody raised in sheep crosslinked to protein G Dynabeads using BS3 (ThermoFisher Scientific). RNMT complexes bound to beads were washed with 100 mM triethylammonium bicarbonate (TEAB) and reduced by incubation in 10 mM DTT/100 mM TAB for 30 minutes at 50°C. Protein alklyation was performed using freshly made 100 mM IAA (iodoacetamide) in 100 mM TEAB. This was followed by an overnight incubation with 1 μg of Trypsin (ThermoFisher Pierce) to allow for protein digestion on-beads. Digested peptides were purified using SP3 beads (Cytiva) and eluted in dH2O. Samples were analysed using ITMS (Ion trap MS) and HCD (Higher-energy collisional dissociation). Data were analysed using CWF Basic and PWF tribrid SequestHT percolator fusion HY workflow within Proteome Discoverer (ThermoFisher Scientific, version 2.4.0.305). Proteins were then filtered using a set of filters on the Proteome Discoverer (ThermoFisher Scientific). For phosphorylation modification detection, data were analysed using CWF Basic and PWF Fusion Basic SequestHT workflow in Proteome Discoverer version 2.2.0.388. To detect serine/threonine phosphorylation modifications, the dynamic modifications were configured to Phospho / +79.966 Da (S, T).

For recombinant RNMT mass spectrometry experiment, *in vitro* phosphorylated RNMT was resolved onto gradient SDS-PAGE gel (4-12%). After Coomassie staining using Instantblue (Abcam), bands were excised prior to in-gel tryptic digestion. Protein bands were cut and transferred to a micro-centrifuge tube. Excised gels were further cut into smaller pieces. The gel pieces were washed for 15 minutes sequentially with 200 μL of Milli-Q water, 200 μL of acetonitrile, 200 μL of 100 mM triethylammonium bicarbonate (TEAB), and 200 μL 100 mM TEAB/acetonitrile (50:50 v/v). The pieces were then washed with 100 μL of acetonitrile for 10 minutes and briefly dried in speed Vac. Reduction of cysteine residues was performed by adding 50 µl of DTT at 20 mM followed by incubation at 56°C for 1 hour. Alkylation was performed by adding of 50 µl of 50 mM iodoacetamide (IAA) prepared in 20 mM TEAB and incubation for 30 minutes at room temperature. Samples were then centrifuged, and the resulting supernatants were discarded. Trypsin was used for in gel digestion of proteins at a final concentration of 12 μg/mL (in 20 mM TEAB) and incubation at 30°C for 16 hours on a shaker. Acetonitrile was added and peptide mixtures were extracted by shaking at 30°C for 15 minutes. The resulting supernatant was transferred to a fresh microcentrifuge tube.

Peptides in the gel pieces were further extracted by adding of 5% formic acid and shaking for 15 minutes followed by adding the same volume of 100% acetonitrile and shaking for another 45 minutes. The supernatant was then collected and transferred to the first fraction of the peptide mix. A final wash of the gel pieces was carried out using acetonitrile for 10 minutes and the resulting supernatant was pooled with the peptide mix. The LC-MS analysis was performed by the FingerPrints Proteomics Facility at the University of Dundee. Analysis of peptide readout was performed on a Q Exactive™ plus, Mass Spectrometer (ThermoFisher Scientific) coupled with a Dionex Ultimate 3000 RS (ThermoFisher Scientific). The data were analysed using the ptmRS module within Proteome Discoverer (ThermoFisher Scientific, version 2.2).

For RNMT immunoprecipitation in primary murine neurons, IPs were carried out as previously described. Eluted material was resolved on 4-12% NuPAGE Bis-Tris gel using MOPS buffer and the gel was stained with Coomassie Instantblue (Abcam). Excised gel fragments were processed as described above. The LC-MS analysis was performed by the FingerPrints Proteomics Facility at the University of Dundee. The trypsin-digested peptides were run on a Q-Exactive HF (ThermoFisher Scientific) instrument coupled to a Dionex Ultimate 3000 HPLC system (ThermoFisher Scientific). The raw data were analysed in MaxQuant version 1.6.2.10 using label-free analysis.

### Immunofluorescence staining

Immunofluorescence staining was performed as in^20^. Briefly, PFA-fixed cells and tissue sections were permeabilised with 1% Triton X-100 in PBS and blocked with 10% donkey serum in PBS with 0.05% Tween-20. After an overnight incubation with primary antibodies, slides were washed three times with PBS/0.05% Tween-20 and incubated with appropriate Alexa Fluor conjugated donkey secondary antibodies for 1 hour at room temperature. Stained slides were mounted in DAPI mounting medium (ibidi). Images were captured using confocal microscopy (Leica SP8 and Nikon A1R) and Evos FL AMF4300 system. The following objectives were used for the Leica SP8 system (Model: DMI8-CS): HC PL APO CS 10X/0.40 DRY, HC PL APO CS2 20X/0.75 DRY and HC PL APO CS2 63X/1.40 OIL. For the Nikon A1R point-scanning confocal microscope, images were acquired using a 20X objective (20x/0.75 Plan APO DIC N2). Images and figure panels were processed using OMERO^5^. For RNMT cytoplasmic and nuclear intensities in primary neurons, CellProfiler (Version 4.2.5) software ^6^ was used. For the cytoplasm (neuronal processes), primary objects between 5–20 pixel unit were identified using the red channel (MAP2 staining). For nuclei, primary objects between 15-40 pixel unit were identified in the DAPI channel. The following parameters were used to identify all primary objects: Threshold strategy: Global, Thresholding method: Otsu, Threshold smoothing scale:1.3488, and Threshold correction factor:1. The intensity in the identified primary objects were then measured using the green channel (RNMT staining).

### Size exclusion chromatography (SEC)

Cells and organ tissue were lysed as described above. Lysates were cleared by spinning at 13,000 rpm for 15 minutes and filtered using 0.22-μm Costar Spin X columns (Corning). 0.25 ml of 8 mg/ml of total extract was injected through a 500 μl sample loop and allowed to resolve on Superdex 200 Increase 10/300 GL SEC column (Cytiva) in 50 mM Tris–HCl, pH 7.5, 1 mM EDTA pH 8, 100 mM NaCl, 0.03% Brij-35 (Merck), and 2 mM DTT at 0.5 ml/min flow rate. A total of 1.5 column volume was collected (0.5 ml/fraction). Fractions were boiled in 2x Laemmli buffer and separated onto 4-12% SDS-PAGE gels for western blot analysis. Gel filtration molecular weight markers (MWGF200-12,000-200,000 Da, Sigma) were used to determine the approximate size of proteins.

### Neurite length measurement

The hiPSC derived dopamine neuronal neurite growth assessment was carried out in 24-well plates using the live imaging platform IncuCyte® S3 (Sartorius). Briefly, three embryoid bodies were seeded/ cm^2^, and neuronal differentiation was induced as previously described. The neurites of differentiated neurons were stained using monoclonal mouse anti-Tuj1 (Sigma clone TU-20, 1:250) and Alexa Fluor 594– conjugated donkey anti-mouse secondary antibody. Nuclei were stained with NucSpot 470 (Biotium). Images were captured per well with exposure 300 ms (GFP) and 400 ms (RFP) using the 10X objective. The average neurite length (mm) per nucleus was measured using the combined colour metrics in the IncuCyte Neurotrack analysis module using the following analysis parameters: TOP-HAT, Radius 20 μm, Threshold (RCU) 5, clean-up 0, adjust size 0, min cell width 7 μm, Neurite coarse sensitivity 10, Neurites fine sensitivity 0.65, neurite width 1 μm. The bar graphs indicate the mean ± standard deviation from the whole field images from three independent experiments.

### *In vitro* phosphorylation

Recombinant GST-CaMKII alpha (SinoBiological) was pre-incubated with 1.2 μM calmodulin (Sigma) for 15 minutes at 30°C in kinase buffer (50 mM Tris, pH 7.5, 10 mM MgCl2, 2 mM DTT, 0.1 mM EDTA, 2 mM CaCl2, 200 μM ATP, 0.01% Brij35). 500 nM of pre-activated GST-CaMKII was then incubated with 0.5 μg His tagged RNMT proteins at 30°C for 90 minutes or as indicated in 20 μl of kinase buffer. For ^32^P labelling, 0.2 μl [γ-^32^P] ATP (PerkinElmer) was added. *In vitro* phosphorylation reactions were stopped by either boiling in 2x Laemmli buffer or snap-frozen on dry ice for the RNMT methyltransferase activity assay. Samples were resolved on a 4-12% SDS-PAGE gel and ^32^P-labeled proteins were visualised by autoradiography.

### Animal perfusion and brain sectioning

Male CD-1 mice (25 and 6.5 weeks old) were terminally anaesthetised by administering an intraperitoneal (IP) injection of an overdose of Euthatal solution in PBS (40 mg/Kg,10 ml/Kg). After confirming unresponsiveness, animals were transcardially perfused with 20 ml of PBS followed by 20 ml of 4% paraformaldehyde to allow the rapid fixation of organs. Brains were removed and placed in 30% sucrose/PBS overnight at 4°C. Brains were isolated and sectioned using a freezing sledge microtome at 50 μm thickness. Sections were stained as described in the IF section.

### RNMT methyltransferase (MT) activity assay

An RNMT methyltransferase (MT) assay was performed according to (Varshney et al. 2016). Briefly, 20-30 ng of recombinant human RNMT or immunopurified GFP-RNMT (on-beads) was incubated at 30°C for 10 minutes with 2 µM S-adenosyl methionine and ^32^P -GpppG-capped in vitro transcribed RNA in 10 µl MT buffer (50 mM Tris, pH 8, 6 mM KCl, 1.25 mM MgCl2) and 0.5 µl RNAsin (Promega). RNA was purified by phenol chloroform extraction and precipitated in sodium acetate or ammonium acetate. The RNA pellet was then resuspended in 0.4 µl 50 mM sodium acetate and incubated with 1µl (0.1units) P1 nuclease for 37°C for 15 minutes. 1 µl of reaction was resolved on PEI cellulose in 0.4 M ammonium sulphate. Spots were visualised by phospho-imaging and quantified in the AIDA image analyser.

### RNMT T317 molecular dynamics (MD) simulations

Simulation systems of RNMT in the presence or absence of SAM and the cap (G0pppG1) were prepared as described by Bueren-Calabuig et al.^21^. The positions of the substrates were obtained from coordinates of SAH in the human RNMT-RAM crystal structure (PDB 5E8J) and the cap in the *E. cuniculi* Ecm1 structure (PDB 1RI2)^22,23^. All protein chains were capped using acetyl (ACE) groups on the N-termini and amino (NME) groups on the C-termini. Protonation was performed at neutral pH using the Reduce module^24^. Phosphorylation was performed in PyMOL to prepare the systems containing pT317. Preliminary simulations were performed with phosphorylations in their unprotonated (-2) and protonated (-1) states. Both states showed comparable behaviour (data not shown), therefore, we present only the data from the unprotonated simulations. The systems were prepared with the AMBER16 suite and the LEaP module. Octahedral boxes of TIP3P water were used to solvate the systems, with the box edges being >15 Å from the protein. Na^+^ or Cl^-^counter ions were added to the systems until the charge was neutralised.

All simulations were performed in AMBER16 using the pmemd.cuda module^14^, using the ff14SB force field modified to include the substrate parameters described by Bueren-Calabuig et al. and the phosphoserine/phosphothreonine parameters described by Homeyer et al.^21,25^. A combination of steepest descent and conjugate gradient energy minimisation was performed to minimise the systems before equilibration. The systems were then heated from 100 K to 310 K over 25 ps, with position restraints on the solute. Equilibration of the systems was performed for 200 ps as the solute restraints were decreased. Finally, 20 ps of equilibration were performed without solute position restraints. During heating and equilibration, a 2 fs time step was used. After equilibration, hydrogen mass repartitioning was performed, and the production runs were performed with a 4 fs time step^26^. Constant pressure (1 atmosphere) and temperature (310 K) were imposed with the Berendsen barostat and thermostat^27^. The SHAKE algorithm was used to restrain hydrogen bond lengths and a 10 Å non-bonded cutoff distance used^28^. For all systems, 200 ns of MD was performed in triplicate. A summary of all the simulations described in this work is given in (Figure S7C).

All simulation trajectories were inspected with the VMD and PyMOL packages^13,29^. Analysis of the simulations was performed on trajectories with frames saved at 40 ps intervals primarily using the CPPTRAJ module in the AMBER16 suite^30^. In particular, CPPTRAJ was used to analyse the interatomic distances, angles, and root-mean-square deviations (RMSDs). Electostatic potentials were generated with the Adaptive Poisson-Boltzmann Solver (APBS) methods using the PyMOL APBS plugin ^31^.

### RNMT CLIP

RNMT seCLIP in primary neurons was performed with biological triplicates (CLIP and corresponding SMI (size-matched input) with an additional non-crosslinked IgG control (and corresponding SMI), as previously described (Van Nostrand et al, 2017) with the following alterations. Primary neurons were cultured as previously stated and harvested at DIV 16. To determine experimental conditions for seCLIP experiments, we used previously defined optimal conditions established in pilot experiments performed on mESC (data not shown) for the endogenous RNMT polyclonal sheep antibody. Cells were washed once in ice-cold PBS and subjected to UV-crosslinking at 254 nm, 400 mJ/cm2 (Stratalink). Cells were lysed in ice-cold lysis buffer (50 mM Tris-HCl pH 7.5, 100 mM NaCl, 1% NP40, 0.1% SDS, 0.5% Sodium deoxycholate), supplemented fresh with protease inhibitors, 440U/ml Murine RNase inhibitor (NEB, 40 U/µl) and 40U/ml Turbo DNase (ThermoFisher Scientific, 2 U/µl). Limited RNase digestion was performed using freshly diluted RNase I (Ambion, 100U/ µl) in PBS (1:20 and 1:200 for high and low RNase conditions respectively), for 5 minutes at 37 °C. 6 µg of RNMT antibody or sheep IgG control (ThermoFisher Scientific) was incubated with 100 µl of protein G Dynabeads (Thermofisher Scientific) per sample for 45 minutes with rotation at room temperature. Antibody coupled beads were then incubated with prepared cell lysates overnight at 4 °C. After immunoprecipitation, 20 µl of bead/lysate per sample was removed to serve as an SMI control and another 20 µl removed for imaging purposes in accordance with the seCLIP protocol ^32^. CLIP and input samples were processed as previously described before separation by SDS-PAGE on NuPAGE 4-12% Bis-Tris gels and transferred onto nitrocellulose membranes. Protein-RNA complexes were isolated from nitrocellulose membranes at the appropriate size range guided by QC steps performed prior to library preparation (see radiolabelling method below). Subsequent steps, including preparation of seCLIP and corresponding SMI cDNA libraries, were performed without deviation from the published seCLIP protocol ^32^. The number of PCR cycles used to amplify cDNA libraries were defined using a diagnostic qPCR quantification of cDNA, by taking 3 cycles fewer that the Ct value for each sample as stipulated by the seCLIP method. Following library amplification with the optimal PCR cycle number, library cleanup was performed as per the seCLIP protocol, and cDNA libraries quantified using the Tapestation and Bioanalyzer platforms (Agilent). Samples were sequenced using the NovaSeq 6000 platform (Illumina) with the SP reagent kit (200 cycles, 2x100bp paired end reads) to generate a maximum of 20 million reads per sample.

Radiolabelling of RNMT-bound RNA was performed during our optimisation process to ensure protein-RNA complexes of the correct size range could be observed following IP, serve as an accurate guide when isolating RNMT-RNA complexes from nitrocellulose membranes during the seCLIP library preparation, and to ensure RNase digestion conditions were optimal. After IP, washes and fastAP steps were performed as described for seCLIP library generation, samples were washed 1x in PNK buffer (50 mM Tris-HCl, 10 mM MgCl2, 50 mM NaCl), before resuspension in 20 µl of PNK mix per sample (1 µl PNK (NEB), 0.25 µl 32P-γ-ATP (6000Ci/mmol), 2 µl 10x PNK buffer, 16.6 µl ddH2O) and incubation at 37 °C for 30 minutes. Samples were spun down and 100 µl of phosphatase buffer was added to stop the reaction, beads were then washed twice with 1 ml lysis buffer and once with no-salt wash buffer before resuspension in 20 µl of 1.5x NuPAGE loading buffer. Samples were eluted from the beads at 70 °C for 10 minutes, and the supernatant collected for SDS-PAGE as described for seCLIP. Radiolabelled samples were transferred to nitrocellulose membranes at 30V for 1 hour at 4 °C and exposed overnight before imaging by phospho-imager (Amersham Typhoon, Cytiva).

For the RNA binding analysis of GFP-RNMT WT and T317 mutants in HEK293 cells, IR-CLIP was carried out as described in (Zarnegar et al. 2016, Kaczynski, Hussain and Farkas 2019). 80% confluent HEK293 cells expressing the different GFP-RNMT mutants were UV cross-linked as previously described. Cells were then lysed in (50 mM Tris pH 7.4, 100 mM NaCl, 0.1% SDS, 1% NP40, 0.5% Sodium deoxycholate) supplemented with Protease inhibitor cocktail III (Sigma), Murine RNAse inhibitor (NEB) and Turbo DNase (ThermoFisher Scientific). Lysates were diluted to 1 mg/ml concentration prior to RNAse treatment at 37 °C for 5 minutes with a final concentration of 1:20000 RNase 1. Immunoprecipitations were carried out overnight at 4 °C using 1 µg of polyclonal anti-GFP (Bio-Rad) antibody and 10 µl of protein G Dynabeads (Thermofisher Scientific). An input of 4% of IP/lysate was taken before the IP is washed with low salt wash buffer (20 mM Tris pH 7.4, 10 mM MgCl2, 0.2% Tween) and high salt wash buffer (50 mM Tris pH7.4, 1M NaCl, 0.1% SDS, 1% NP40, 0.5% Sodium deoxycholate). The immunoprecipitated RNA was then dephosphorylated with T4 PNK (Thermofisher Scientific) for 15 minutes at 37 °C. Then, samples were washed twice with low salt wash buffer before the IR adaptor was ligated using T4 RNA ligase 1 (NEB) for 75 minutes at room temperature. IPs were then washed using a salt gradient before being resuspended in 2X LDS buffer with 10% DTT and eluted at 70 °C for 10 minutes. Eluted samples were resolved on 4-12% SDS PAGE gels and transferred to membrane at 30V overnight at 4 °C. The membrane was then imaged using the LICOR imaging system in the 800 channel, blocked and processed for western blotting for proteins of interest.

### SeCLIP data analysis

After standard demultiplexing, paired-end reads were subjected to several pre-processing steps. Data were subject to initial QC using fastQC and reads extracted using umi-tools in extract mode. Adapters were trimmed from both reads using cutadapt (version 1.18, run with python 2.7.15) with the following options: -a AGATCGGAAGAGCACACGTCTGAACTCCAGTCAC -AAGATCGGAAGAGCGTCGTGTAGGGAAAGAGTGT --trim-n -m10. Trimmed reads were aligned to the mm10 version of the mouse genome, using STAR (Dobin et al. 2013), version 2.5.2b) and an index generated with the option: --sjdbOverhang 99 (corresponding to 100bp sequenced reads). Aligned reads were demultiplexed using umi-tools in dedup mode to resolve PCR duplicates in the data. Demultiplexed reads were then sorted and indexed using SAMtools ^10^ before being used in downstream analyses. Technical replicates were merged using SAMtools and processed using various Bedtools commands. To obtain crosslink sites, merged .bam files were converted to .bed files before shifting each coordinate by 1nt to account for stalling of the reverse transcriptase at the site of UV-crosslinking. After inferring the genome coverage, each sample was converted to bedgraph format for visual inspection in the UCSC genome browser. For the crosslink reads, count for each transcript, individual BAM files were processed using iCOUNT package ^9^.

The DA neuronal RNA-seq data and seCLIP data correlation analysis was performed first by removing transcripts with read counts less than 10-fold enrichment over IgG control CLIP. The two datasets were then combined using a mouse/human homologue database from BioMart to find all matched genes across the two species. The correlation plots were generated using the top 10% translated neuronal transcripts identified in ^33^. Crosslink profiles along RNA-seq related transcripts were achieved using deepTools plot profiles ^11^. For the genomic feature analysis of the RNMT-bound transcripts, the top 10% targets identified in RNMT seCLIP in primary neurons were used. The coordinates of genomic features (CDS, 3′ UTR, 5′ UTR) were obtained from Ensembl Biomart gene feature database.

### RNA-seq

Cytoplasmic RNA was extracted from hiPSC-DA neurons (3 biological replicates per cell line) according to ^34^. Sample QC, ribosomal RNA depletion, RNA library preparation and RNA sequencing were carried out by Genewiz (Azenta Life Sciences, Germany) using Illumina NovaSeq with 150-bp pair-ended reads. Reads quality and suitability for downstream bioinformatics analysis were assessed using FastQC. Reads were aligned to human genome assembly GRCh38/hg38 using STAR ^7^ and transcripts were quantified using featureCounts^35^. Read counts were normalized to the library size and differential expression was analysed using EdgeR exactTest ^8^. Only transcripts expressed above 1 reads per million in 3 replicates were considered for downstream analysis. The differentially expressed transcripts were identified with false discovery rate (FDR)<0.05. Data were plotted in RStudio. To identify RNMT regulated RNAs, a published Ribo_seq dataset in neuropil and somata of rodent hippocampal excitatory neurons was used ^33^. Matching transcripts between hiPSC-DA neurons and the Ribo_seq dataset were assessed for differential gene expression in DA neurons expressing RNMT mutants. The top 10% translated transcripts from the published Ribo_seq dataset based on the mean expression value, were also assessed for the differential gene expression analysis. The Gene Ontology (GO) analysis was performed using WebGestalt ^12^.

### RT-PCR

Cytoplasmic and nuclear RNAs were extracted using Norgen cytoplasmic & nuclear RNA purification Kit (NorgenBioteK). 300 ng of RNA was used in 20 μl cDNA synthesis reactions using the iScript cDNA Synthesis Kit (Bio-Rad). PCR was carried out using EvaGreen supermix (Bio-Rad) on the CFX384 Touch realtime PCR system (Bio-Rad). Ct values of target genes were normalized to the corresponding Ct value of GAPDH. Primer sequences are listed in the Key Resources Table.

